# Suprathreshold contrast response in normal and anomalous trichromats

**DOI:** 10.1101/788331

**Authors:** Kenneth Knoblauch, Brennan Marsh-Armstrong, John S. Werner

**Affiliations:** Univ Lyon, Université Claude Bernard Lyon 1, Inserm, Stem Cell and Brain Research Institute U1208, 69500 Bron, France; National Centre for Optics, Vision and Eye Care, Faculty of Health and Social Sciences, University of South-Eastern Norway, Hasbergsvei 36, 3616 Kongsberg, Norway; Vision Science and Advanced Retinal Imaging Laboratory, Department of Ophthalmology and Vision Science, University of California, Davis, USA

## Abstract

Maximum Likelihood Difference Scaling was used to measure suprathreshold contrast response difference scales for low-frequency Gabor patterns modulated along luminance and L-M color directions in normal, protanomalous, and deuteranomalous observers. Based on a signal-detection model, perceptual scale values, parameterized as *d′*, were estimated by maximum likelihood. The difference scales were well fit by a Michaelis-Menten model, permitting estimates of response and contrast gain parameters for each subject. Anomalous observers showed no significant differences in response or contrast gain from normal observers for luminance contrast. For chromatic modulations, however, anomalous observers displayed higher contrast and lower response gain compared to normal observers. These effects cannot be explained by simple pigment shift models and support a compensation mechanism to optimize the mapping of the input contrast range to the neural response range. A linear relation between response and contrast gain suggests a neural trade-off between them.

## 1. Introduction

Anomalous trichromacy is classically defined by abnormal shifts in the mixture of reddish and greenish primaries in metameric matches to a yellowish standard [1]. Observers termed protanomalous use more of the reddish primary in the match, and deuteranomalous observers use more of the greenish one. Based on colorimetric [2–5] and genetic [6, 7] studies, it is generally held that the change in matching behavior is explained by shifts in the peaks between the spectral sensitivities of the middle- (M-) or long- (L-) wavelength sensitive cone photoreceptors compared to normal (Fig. 1a). Specifically, deuteranomaly is thought to arise from a substitution of the normal M-cone with a variant L-cone shifted toward longer wavelengths than the normal M-cone [7, 8] and is denoted by L’. Similarly, in protanomaly the normal L-cone photopigment is replaced by a variant M-cone that is shifted toward shorter wavelengths than the normal L-cone and is denoted by M’.

**Fig. 1.**
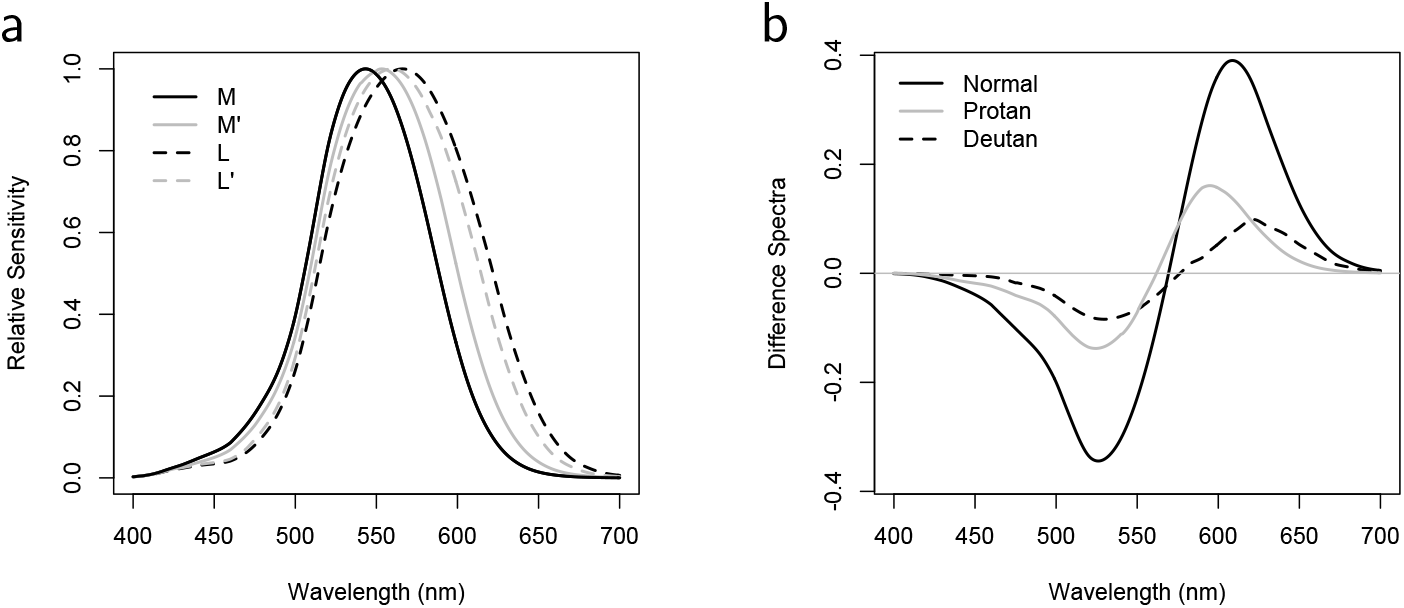
a. DeMarco, Pokorny and Smith fundamentals [12] for normal M and L cones (black solid and dashed, respectively) and average anomalous observers’ M’ and L’ cone spectral sensitivities (grey solid and dashed, respectively). b. Difference spectra of two long-wavelength sensitive cone spectral sensitivity functions for average Normal (black solid, L - M), Protanomalous (grey solid, M’ - M) and Deuteranomalous (black dashed, L - L’) observers. The weights were chosen so that their magnitudes sum to unity and that the net response to an equal-energy light is zero.

Losses in discrimination that many (but not all [9, 10]) anomalous observers display are attributed to the reduction in the spectral signal from the greater overlap of the cone spectral sensitivities. While there is variation in peak separations for both normal [11] and anomalous observers [10], the loss of chromatic sensitivity for average anomalous observers can be visualized by plotting the difference in the two long-wavelength spectral sensitivities, shown in Fig. 1b [12]. The long-wavelength chromatic difference signal is reduced in anomalous observers with respect to the normal curve. The peak-to-trough difference of the protanomalous curve is 41% of the normal, and that of the deuteranomalous, 25% of the normal.

### 1.1. Null model for contrast perception in anomalous trichromacy

Despite affecting approximately 6% of Caucasian males, little is known about the consequences of color anomaly on color appearance [13]. A simple null hypothesis is that the reduced chromatic signal only attentuates the effective contrast of the input to a chromatic differencing mechanism by a factor *α*. We can simulate this using a Michaelis-Menten function as a model for the chromatic response, *R*, as function of contrast, *c*.

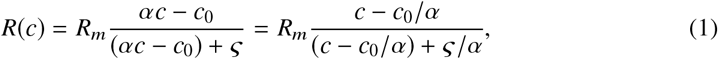

where *R_m_* is the maximum contrast response or response gain, *c*_0_ is a threshold value for perceived contrast and *ς* the contrast at which the response is half-maximal. The inverse of *ς* is sometimes taken as a measure of contrast gain. Dividing through the numerator and denominator by *α* in the right-hand term demonstrates that reducing the effective contrast will modify both the threshold perceived contrast and the semi-saturation constant but will not influence the response gain. Since in anomalous vision it is expected that *α* < 1, the effect will be to increase both the threshold perceived contrast and the semi-saturation constant, i.e., decrease the contrast gain. This is demonstrated in Fig. 2a in which *α* was assigned values of 1.0 (normal) and 0.41 and 0.25, to approximate average contrast losses in protan and deutan anomalous observers, respectively. The protan and deutan curves are shifted to the right with respect to the normal curve and appear to rise more slowly. However, when these curves are plotted on a log contrast axis in Fig. 2b, the multiplicative scaling effects become simple translations along the abscissa with no change in curve shape.

**Fig. 2.**
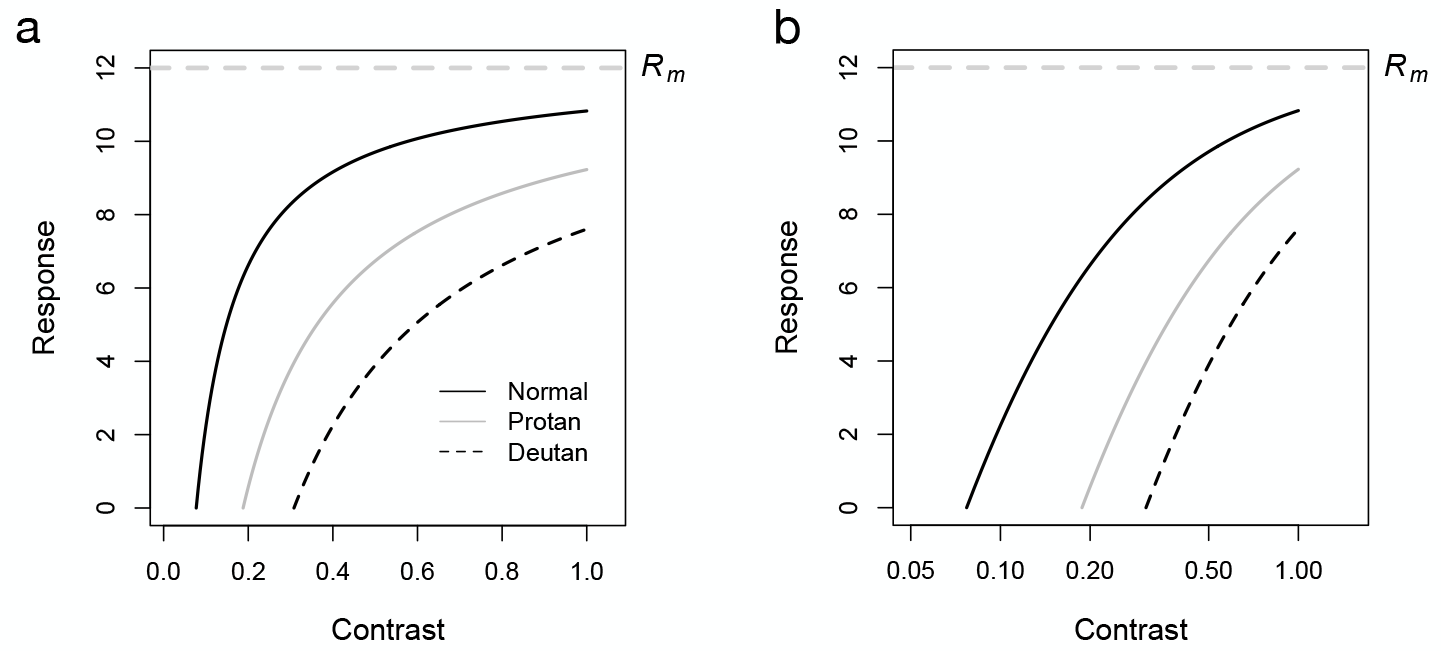
a. Simulated chromatic contrast response curves for Normal, Protanomalous and Deuteranomalous observers based on the assumption that the effective chromatic contrast is reduced in anomalous observers. b. The same three curves plotted on a logarithmically spaced contrast axis.

### 1.2. Previous studies

In a study of hue cancellation, anomalous trichromats showed a relative reduction of chromatic valence along the red-green axis that was correlated with anomaloscope matching range [14]. Several studies in which perceived color differences were estimated by multidimensional scaling (MDS) reported compression of perceived differences along the red-green axis [15–19]. In a study of Visual Evoked Potentials (VEP), several anomalous observers were examined [20]. The study focussed primarily on latencies rather than amplitudes but reported that anomalous observers showed consistently longer latencies along an L-M axis. The evidence from these approaches appears consistent with the reduced chromatic input signal from the greater overlap of cone spectral sensitivities.

Boehm, MacLeod and Bosten [21], however, found that the compression in the perception of large color differences in MDS was much less than that obtained in discrimination experiments. As the discrimination results were well explained by the loss of spectral information at the input, they suggested that a post-receptoral gain amplification occurs in anomalous observers at suprathreshold levels. Support for post-receptoral gain amplification in anomalous trichromacy has also been reported from functional cerebral imaging [22]. These results fit with a theory proposed by MacLeod that the reduced input signal might be mapped onto the full range of neural response, thereby compensating for the genetic effects on the spectral tuning of photopigment spectra [23, 24].

MacLeod described the consequences of two hypotheses to explain possible adaptations of anomalous observers to their reduced spectral information [23]. If anomalous vision was limited by noise at the input (e.g., at the photoreceptors), then such observers would experience poor discrimination at low chromatic contrasts but good discrimination at high chromatic contrasts. On the other hand, if discrimination was limited by output noise at a post-receptoral stage (e.g., at the color-opponent channels), then discrimination would be enhanced at low chromatic contrasts by increasing the channel gain but at the cost of poorer discrimination for large chromatic contrasts. In fact, Boehm et al. interpret their MDS findings as support for increased gain in post-receptoral channels in anomalous trichromacy. MacLeod [23] cites the well documented evidence of a lack of correlation between midpoint match shifts and ranges of Rayleigh matches [9] as circumstantial evidence to support the latter hypothesis as well as discrimination experiments on his own deuteranomalous vision.

Here, we used Maximum Likelihood Difference Scaling (MLDS), a recently introduced scaling method based on a signal detection model [25–27], to compare the change in appearance of suprathreshold contrasts along luminance and L-M chromatic directions in normal and anomalous observers. The method allows contrast response to be evaluated over nearly the entire range of suprathreshold contrasts, thus, allowing the comparison of normal and anomalous responses at extreme contrast levels.

## 2. Methods

### 2.1. Observers

Twenty-seven volunteers (ages 18–49) completed testing. They were recruited through flyers and an online portal. Procedures were explained before any testing, and observers provided written informed consent using a protocol approved by the UC Davis Institutional Review Board. All observers were compensated for their participation.

Equal numbers of participants were classified as normal (mean age = 26.0 yrs), deuteranomalous (mean age = 26.7 yrs) or protanomalous (mean age = 28.9 yrs) trichromats. All participants were male except for two normal-group females. All observers had normal visual acuity (best corrected to 6/6 or better) and had a negative history of retinal disease and neurological disorders affecting vision.

Color vision classification was based on Rayleigh matches with a Neitz OT anomaloscope and the Cambridge Colour Test (CCT) administered in trivector mode using a calibrated monitor (Eizo FlexScan T566). Observers with anomaloscope coefficients between 0.766 and 1.333 were classified as normal. Deuteranomalous observers were identified as those individuals with values above this range and protanomalous below. On the CCT, all participants classified as normal had deutan and protan vector lengths < 100 with mean values of 56 in both cases. For subjects classified as deuteranomalous, the deutan and protan vector means were 532 and 296, respectively. For subjects classified as protanomalous, the deutan and protan vector means were 315 and 680, respectively. This comports with the criteria for classification of normal and anomalous for the CCT. Additional confirmation of classifications was based on the F2 Plate test, the Farnsworth Panel D-15 test, and the American Optical Hardy-Rand-Ritter pseudoisochromatic plates, all administered under a lamp equivalent to illuminant C.

### 2.2. Apparatus, Stimuli and Calibrations

Stimuli were displayed on an Eizo (FlexScan T566) CRT monitor with a 40.3 cm diagonal screen size operating at a resolution of 1280 × 1024 pixels viewed at a distance of 150 cm. The visual path from viewer to stimuli was surrounded by black light baffles internally coated by non-reflective fabric. Observers were refracted for the test distance using standard trial lenses rather than their habitual spectacles if they were tinted or had anti-reflective coating.

Stimuli were displayed with 10–bit color resolution using custom code written in Python 2.7 utilizing the PsychoPy3 package [28] and integrated development environment. Responses were recorded with a 2–button Bluetooth response pad. The display monitor was gamma corrected and chromatically calibrated using a SpectraScan Spectroradiometer 670 placed 150 cm from the screen and the PsychoPy3 IDE’s Monitor Center automated screen calibration tool.

The stimuli were horizontal Gabor patterns defined by the equation

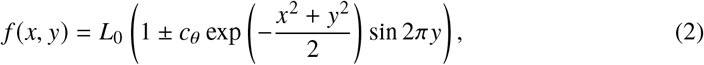

where *L*_0_ is the mean luminance of the screen, *x, y* position in degrees, and *c_θ_* the contrast of the carrier signal along the axis *θ* in color space. The sign of the Gabor term was chosen randomly across trials to generate stimuli varying in phase by 180° in order to minimize local adaptation and afterimages. The stimuli were truncated at 4° diameter (4*σ*) and were in sine phase so that both the space average luminance and chromaticity did not vary across the stimulus. The carrier spatial frequency was 1 cycle per degree (c/deg) and the pattern was offset from a cross-shaped fixation point. Modulation of Gabor patterns was along a luminance axis ([90,0,1], [elevation, azimuth, maximum contrast] in DKL space) or an L-M axis in the isoluminant plane ([0,0,1] in DKL color space) [29]. A brief warning tone preceded each stimulus presentation of 500 ms duration. The steady background was achromatic (CIE (*x, y*) = (0.33,0.35); *Y* = 48.1 cd/m^2^) and continuously present. Using the DeMarco, Pokorny and Smith cone spectral sensitivities for average observers with each color vision type [12], the maximum L-M cone contrasts that could be displayed were estimated as: Normal 0.142, Protanomalous 0.037, and Deuteranomalous 0.041. The CIE (*x, y*) coordinates of these extreme values along the L-M axis were calculated as (0.310, 0.484) and (0.342, 0.226).

### 2.3. Procedures

#### 2.3.1. Preliminary Measurements of Contrast Threshold

Preliminary testing of each individual was conducted to determine a minimum perceived contrast (*c*_0_) for both achromatic luminance modulation and L-M isoluminant stimuli. This was estimated using a two-alternative forced-choice staircase method. Each axis was measured separately with a 5–min dark adaptation period followed by a 3–min re-adaptation period viewing the achromatic background. The Gabor patterns were presented 2.8° above a central 0.5° × 0.5° black fixation cross, and the observer’s task was to press one of two buttons to indicate whether the stimulus was detected or not. The contrast of the Gabor patterns was varied in seven logarithmic steps from 0.125 to 0.875 of the monitor’s maximum displayable (or nominal) contrast in DKL colorspace. This was followed by three more series of threshold tests, each with progressively smaller steps to bracket the previous threshold contrasts. The value of *c*_0_ was taken as 1.7 times the minimum contrast detected in the final subset of 28 stimuli with an error of ± 0.016 contrast units.

#### 2.3.2. Suprathreshold Contrast Response Difference Scaling

To compare contrast response of normal and anomalous trichromats, we used MLDS to estimate suprathreshold Contrast Response Difference Scales (CRDS) [25–27]. Gabor patterns, as described above, were modulated along luminance and L-M color directions in DKL color space on a steady achromatic background in an otherwise dark room. On each trial, 3 Gabor patterns ordered in contrast were presented (0.5 sec, 2.8° eccentricity). The middle value contrast was presented 2.8° above the central fixation cross with the other two straddling the same radial distance to the left and right below the fixation cross (Fig. 3). The lateral positions of the lowest and highest contrast were randomized across trials. A forced-choice response indicated whether the upper pattern was judged as more similar in contrast to the lower left or right pattern. The next trial commenced 0.3 sec after the subject’s response.

**Fig. 3.**
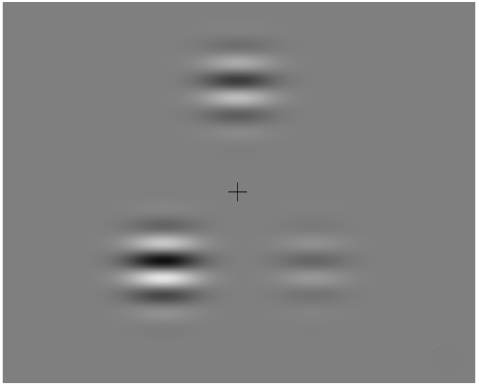
Three luminance Gabor stimuli from an example difference scaling trial. The observer judges which of the two bottom stimuli appears most similar in contrast to the top stimulus.

For each axis of modulation, a range of 9 contrasts was generated in which the lowest was the previously estimated *c*_0_ value of the observer, the highest was 90% of the maximum nominal display contrast, and the sequence of contrasts was equally spaced between these extreme values on a logarithmic axis. Observers were able to order the 9 stimuli by contrast. This resulted in a session of 9!/(3! × 6!) = 84 triads for the forced-choice trials, each preceded by an auditory tone to denote stimulus onset. The order of the contrast triads across trials was random. Each series of 84 trials was completed six times for each axis of stimulus modulation. Rest periods were provided after each series of trials or when a subject requested a break. The rest periods were followed by a 3-min adaptation period to the achromatic background. The two color axes were tested separately. Test runs were distributed over 4 sessions.

### 2.4. Data Analysis and Modeling

#### 2.4.1. MLDS

All statistical analyses were performed using the OpenSource software R [30]. The signal detection model underlying MLDS and the fitting procedure have been previously described [25–27, 31, 32]. In summary, given a set of *p* stimuli ordered along a physical continuum, triples ornon-overlapping quadruples are sampled on each trial. As we used the method of triads, we will develop the model accordingly. Given a trial with physical triples *ϕ*(*a*) < *ϕ*(*b*) < *ϕ*(*c*), we assume a mapping (not necessarily monotonic) onto internal responses, *ψ*(*a*), *ψ*(*b*), *ψ*(*c*). The observer considers the noise-perturbed internal decision variable

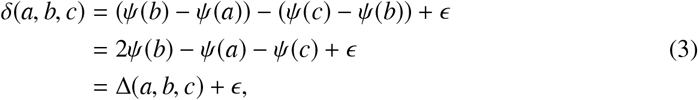

where we have abbreviated *ϕ*(*a*) by *a*, etc. and *ϵ* ~ *N*(0, *σ*^2^). The noise perturbation is called the judgment noise and provides for inconsistencies in the observer’s responses when the decision variable is sufficiently small. The variance of the noise is assumed constant for all triads. If on a given trial *δ* < 0, the observer chooses *a*, otherwise *c*. We code the observer’s responses, *R*, by 1 or 0 depending on whether the choice is stimulus *a* or *c*. From the ensemble of responses to all triads, we compute the log likelihood function assuming that the response on each trial is a Bernoulli distributed variable

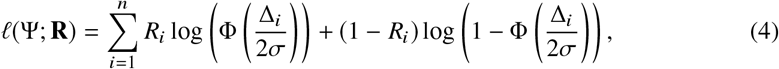

where Φ is the cumulative distribution function for the standard Gaussian, **R** is the vector of responses to all triads and Ψ is the vector of scale values, *ψ_i_*. The scale values and judgment noise are chosen as the values that maximize Eq. 4. While it appears that the model requires estimation of *p* + 1 parameters, to obtain an identifiable solution, we fix the lowest value at 0 and *σ* = 1. This yields *p* − 1 scale parameters to estimate corresponding to *ψ*_2_, *ψ*_3_, …, *ψ_p_*. The parameterization in Eq. 4 renders the scale values in terms of the signal detection parameter *d′* since one unit on the response axis corresponds to the standard deviation of the judgment noise. In practice, we fit the data using functions from the R package **MLDS** [26, 27]. These functions implement the fitting procedure in terms of a generalized linear model with a binomial family and a probit link function [26, 27, 31–33]. The obtained scales are unique up to addition of a constant or multiplication by a coefficient. They have the property that stimulus pairs, separated by equal differences on the ordinate, should appear equally different.

#### 2.4.2. Fitting the response scales

The maximum-likelihood scale values obtained for each observer using Eq. 4, i.e., individual CRDSs, showed a nonlinear dependence on stimulus contrast that was found to be well fit by a Michaelis-Menten function offset by the estimated value of *c*_0_

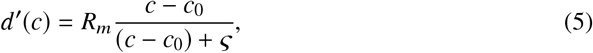

where the other parameters are defined as in Eq. 1. As shown in Fig. 2, the effect of the term *c*_0_ is to shift the function along the contrast axis. As a result, the value of *ς* is determined by the difference from the value of *c*_0_, as shown by the dashed curve in Fig. 4. However, as demonstrated above, *c*_0_ and *ς* are influenced by the effective contrast reduction due to separation of the cone spectra, leading to scale changes on a linear axis, but shape-invariant translations on a log contrast abscissa. Therefore, to assess shape changes that would depend on contrast gain, we examined the value of log_*e*_(*c*_0_/(*c*_0_ + *ς*)), which on the log contrast scale is the difference between the log contrasts at *c*_0_ and the semi-saturation contrast (Fig. 4b).

**Fig. 4.**
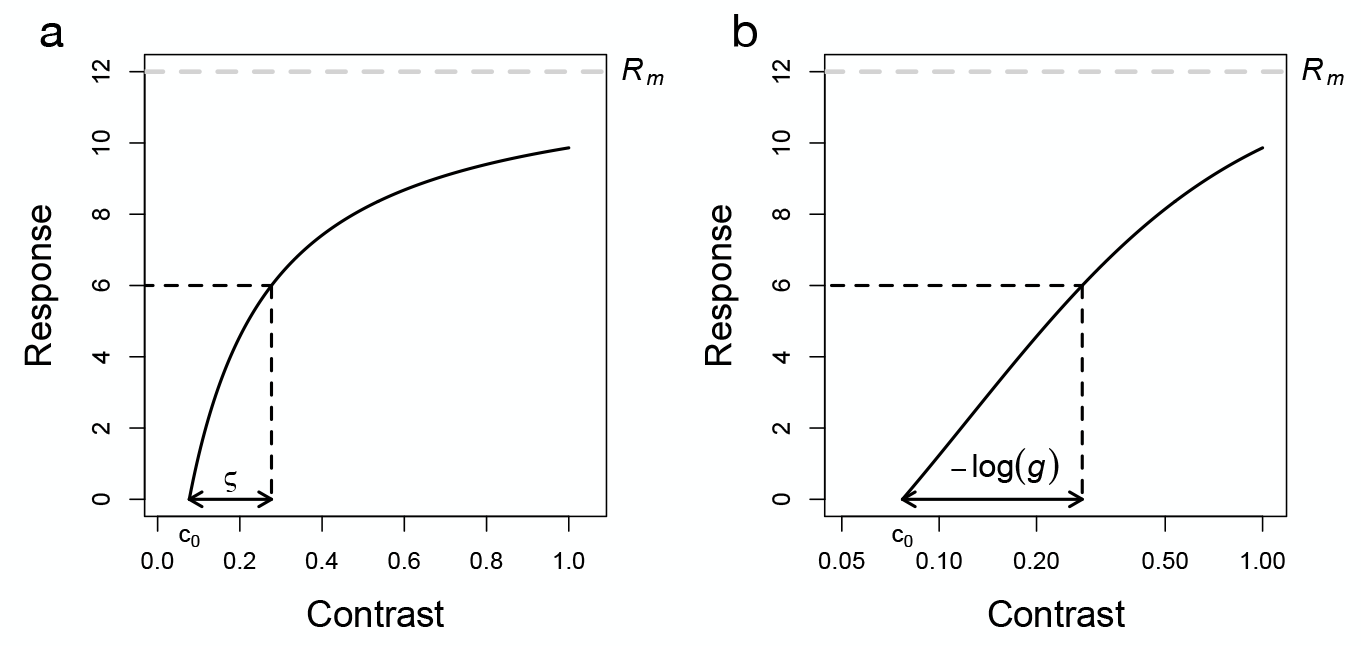
a. Michaelis-Menten function plotted as a function of contrast with the initial value at *c*_0_. The value of *ς* is estimated with respect to *c*_0_. The maximum asymptotic value is shown by the grey dashed line. b. The same Michaelis-Menten function as in a, plotted as a function of the log contrast. The log of the contrast gain, – log(*g*) is defined as the difference between the log contrast at the minimum contrast and that at the semi-saturation constant.

With the aim of estimating this parameter directly, Eq. 5 was reparameterized by solving *g* = *c*_0_/(*c*_0_ + *ς*) for *ς* and substituting giving

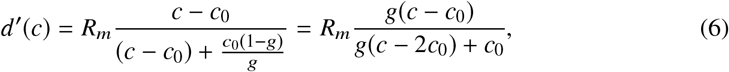

after multiplying the numerator and denominator by *g* and collecting terms in the denominator. By defining *g′* = log_*e*_(*g*), we substituted exp(*g′*) for g to estimate directly the parameter of interest. Such reparameterizations have no effect on the maximum likelihood fit [34].

To compare groups, we fit the data from the three groups of observers with Eq. 6 using a nonlinear mixed-effects model [35].

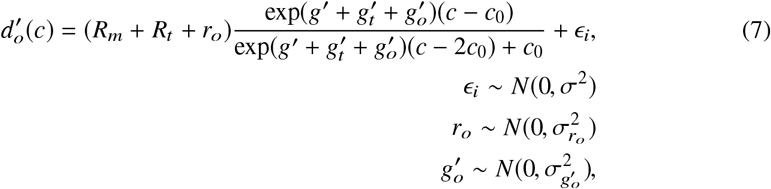

where *R_m_* is the response gain for the normal group, *R_t_* the difference from normal of response gain for the protanomalous or deuteranomalous group, *g′* the log contrast gain for the normal group, and 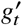, the difference from normal for the protanomalous or deuteranomalous group. The model includes two random effects in addition to the independently and identically distributed mean-zero standard Gaussian random variation across scale values, *ϵ*: random effects of response gain across observer *r_o_* and of the log contrast gain across observer, 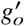, each assumed to be mean-zero Gaussian distributed with variances 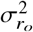 and 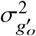, respectively. The difference of response and contrast gain of the anomalous observers from normal was evaluated by assessing whether the terms *R_t_* and 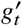, differed from zero. This was accomplished by evaluating Eq. 7 with respect to nested models in which these terms were fixed at zero, using a likelihood ratio test. Data obtained along luminance and L-M axes were fit in separate analyses.

## 3. Results

Figure 5 shows estimated CRDSs for example individual normal (a, d), protanomalous (b, e) and deuteranomalous (c, f) observers. The results from observers classified with the same type of color vision were qualitatively similar and are displayed in the Appendix (Fig. A1) and Supplementary Figs. S1–3 (all supplementary figures and tables referred to in this article can be downloaded from https://www.sbri.fr/sites/default/files/supplementary.pdf). Response scales for luminance contrast stimuli are indicated as filled points and L-M as unfilled points in Fig. 5. The points indicate averages of data collected from sessions on the same day. The curves are the nonlinear least-squares Michaelis-Menten fits which describe the data well in all conditions. The estimated parameters to reproduce each of the individual curves are provided in Supplementary Tabs. S1–2, for data collected for contrasts along the luminance and L-M axes, respectively. Standard errors indicated below and in the Supplementary tables were obtained from the variance-covariance matrix at the maximum likelihood. In order to perform the fits, the data from different days were shifted vertically to minimize the vertical distances between data sets, as such a transformation has no effect on the predicted responses. The shifts required were always small, less than 1% of the response range. On the top row, the data are plotted in nominal contrast with a value of 1.0 corresponding to the maximum output of the display. The estimated minimal contrasts, *c*_0_, for the two anomalous observers are slightly higher than the value for the normal along the luminance axis (N: 0.008; P: 0.016; D: 0.023), but much higher than the normal along the L-M axis (N: 0.037; P: 0.121; D: 0.351). The estimated luminance *R_m_* values (± 1 standard error) were similar for the three observers (N: 10.5 ± 0.53; P: 10.4 ± 0.35, D: 11.9 ± 0.49), but the L-M values were systematically lower for the anomalous observers (N: 6.9 ± 0.42; P: 4.7 ± 0.14; D: 3.4 ± 0.16).

**Fig. 5.**
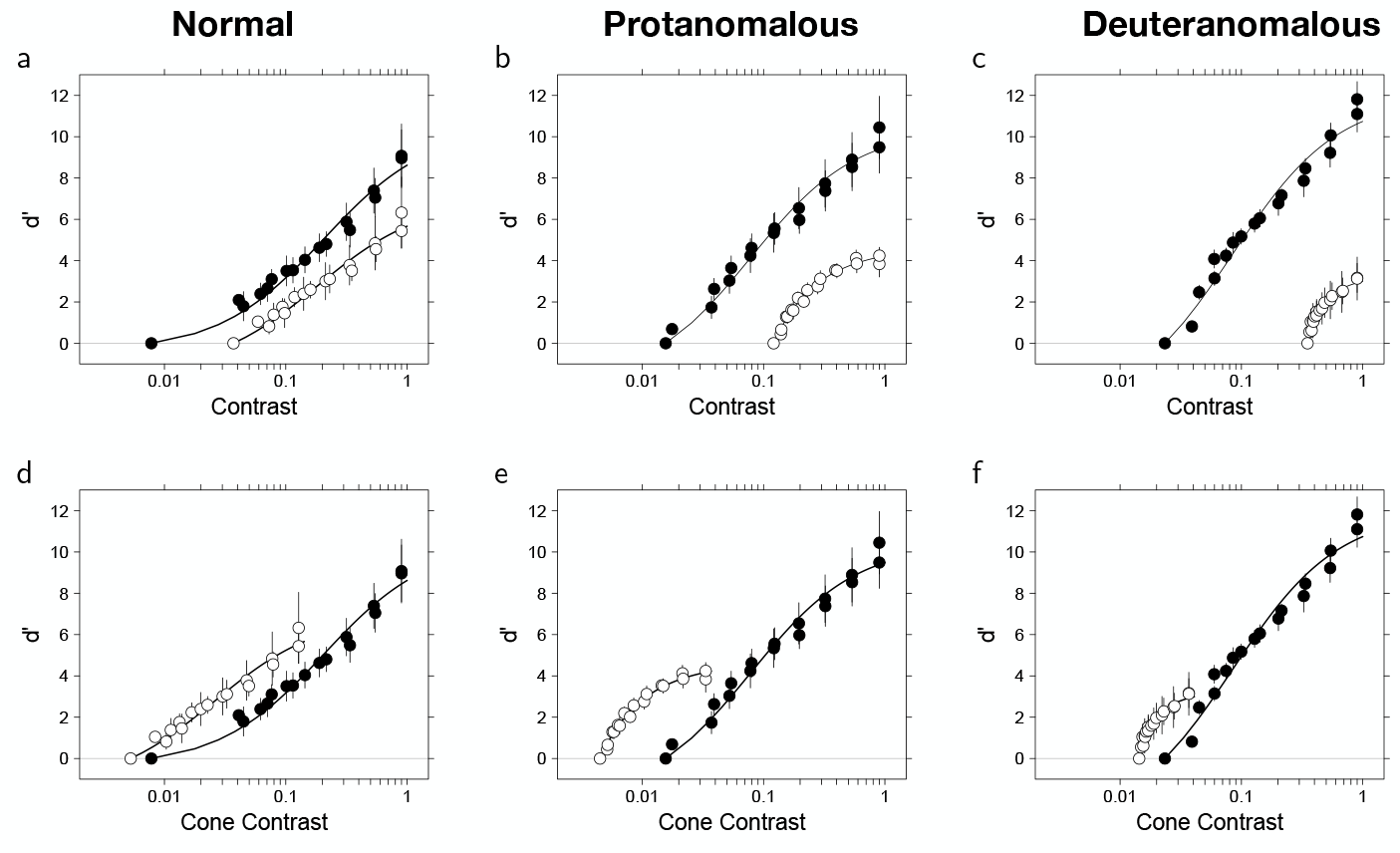
CRDSs estimated by MLDS measured along the luminance (filled symbols) and L-M (unfilled symbols) axes in color space. The top row (a, b, c) shows data from individual Normal (N1), Protanomalous (P4) and Deuteranomalous (D1) observers plotted in nominal contrast units. The bottom row (d, e, f) shows the same data from the respective observers replotted as a function of cone contrast. The abscissa values are logarithmically spaced on all graphs. The solid curves are Michaelis-Menton functions best-fit by nonlinear least squares. The error bars are 95% confidence intervals.

For anomalous observers, the initial branch of the curves appears to rise more steeply to an asymptotic value than for the normal observers along the L-M axis as shown by the estimates of *g′* (N: −1.93 ± 0.127; P: −0.59 ± 0.039; D: −0.24 ± 0.028). It is less obvious from the graphs but for these two observers *g′* is also higher along the luminance axis (N: −3.34 ± 0.122; P: −1.95 ± 0.094; D: −1.73 ± 0.104). The maximum obtainable L-M cone values for an average observer of each color vision type (see Sect. 2.2) were used to rescale the L-M contrasts in the bottom row of graphs. In cone contrast units, all observers responded to the L-M gratings at lower cone contrasts than the luminance gratings [36]. However, with the adjustment to cone contrasts the *c*_0_ values along the L-M axis for all three groups of observers converged toward similar values.

### 3.1. Minimal perceived contrast estimates

Figure 6a shows the *c*_0_ values in nominal contrast for the three classes of observer along both axes tested. To homogenize the variance, a 2-way analysis of variance (ANOVA) was performed on log(*c*_0_), with factors color vision type and color axis tested. The interaction was significant (F(4,48) = 8.175, p ≪ 0.001) indicating that the relative dependence of *c*_0_ on color vision type differed between the two axes tested (Supplementary Tab. S3). Examination of the model coefficients provided no evidence of differences between anomalous and normal observers along the luminance axis (P vs N: t(48) = 0.22, p = 0.83; D vs N: t(48) = −0.09, p = 0.93) but strong evidence for a difference along the L-M axis (P vs N: t(48) = 4.4, p ≪ 0.001; D vs N: t(48) = 5.37, p ≪ 0.001) (Supplementary Tab. S4). The open triangles plotted with the anomalous data in the L-M plot indicate the expected reduction in sensitivity with respect to the mean normal *c*_0_ value due to the relative peak-to-trough reduction of the L-M functions because of the reduced spectral separation of the photopigments (Fig. 1b).

**Fig. 6.**
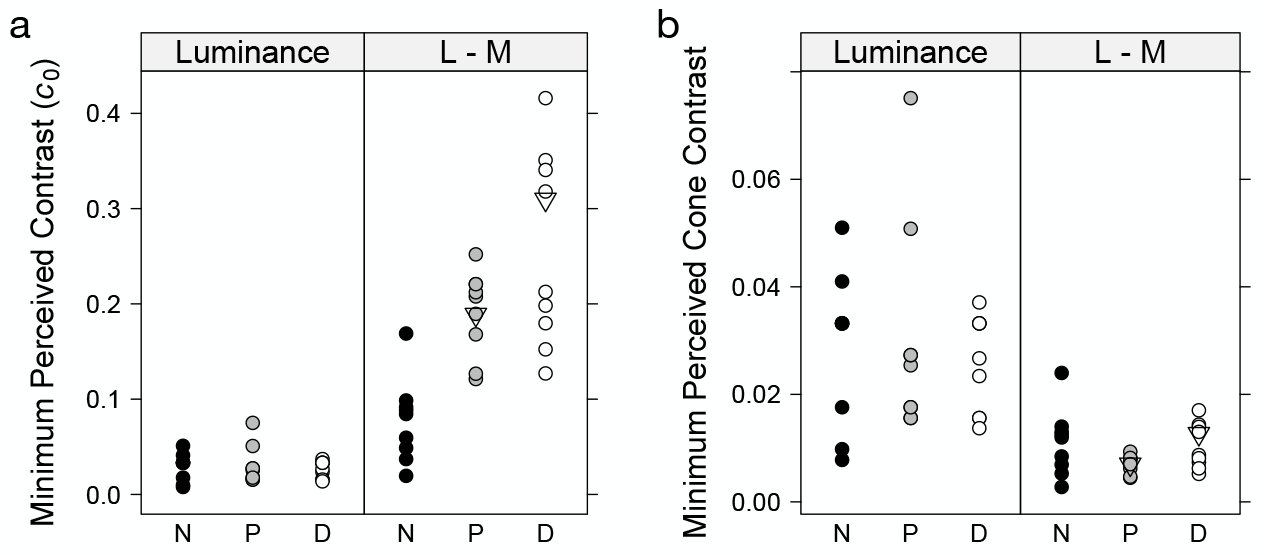
a. Individual estimates of the minimally perceived contrast, *c*_0_ in nominal contrast units for Normal (N), Protanomalous (P) and Deuteranomalous (D) observers along luminance and L-M axes. The open triangles on the L-M plot for protan and deutan observers correspond to the increase of the mean normal *c*_0_ value expected from the reduced separation of the cone spectral sensitivities. b. Individual estimates of *c*_0_ adjusted in units of cone contrast for the 3 classes of observer and both axes of color space.

When the *c*_0_ values are expressed as cone contrasts (Fig. 6b), however, the L-M values no longer differ among groups as indicated by a one-way ANOVA (F(2, 24) =1.5, p = 0.24) (Supplementary Tab. S5). We did not re-analyze the luminance values because these are unchanged in terms of cone contrasts. Note, however, that the chromatic values are lower than the luminance values. The results support the hypothesis that at the level of cone contrasts, all observers require similar neural signals to perceive a minimal chromatic contrast [37]. Equation 1 suggests that uncertainty in *c*_0_ could be attributed, at least in part, to individual variability in photopigment spectra.

### 3.2. Group analyses of contrast response

We fit the Michaelis-Menten function to the data of all observers using a nonlinear mixed-effects model [35], as specified in Eq. 7. Equation 6 was used as the fixed-effect component. An observer-dependent random effect was attributed to both *R_m_* and *g′*, each assumed to be normally distributed with its own variance term. The constant *c*_0_ was not estimated, and the individual values were used. Analyzing the data from luminance and L-M directions together revealed inhomogeneity in variance across the two conditions. Therefore, the data from each axis were analyzed separately.

The most complex models fit included different variance components for each color vision type (Supplementary Tabs. S6–7 and S20–21). Simpler models in which the three color vision types shared a common variance term were then fit (Supplementary Tabs. S8–9 and S22–23). Likelihood ratio tests did not support significant differences between the models (Luminance: *χ*^2^(18) = 7.54, p = 0.99; L - M: *χ*^2^(18) = 18.78, p = 0.41) (Supplementary Tabs. S10 and S24), so we continue with the simpler models.

Figure 7a shows the fixed-effect or population estimates for the CRDSs for the luminance axis for the three observer classes. The upper asymptote is similar for normal and protanomalous observers but is slightly elevated for the deuteranomalous (Fig. 8a, Supplementary Tab. S8). A likelihood ratio test, however, comparing models in which *R_m_* could vary among the three groups with the nested model in which *R_m_* was constrained to be the same across groups, yielded no evidence for a significant difference (*χ*^2^(2) = 1.23, *p* = 0.54) (Supplementary Tab. S13). The anomalous curves appear to rise a little more steeply, suggesting a higher contrast gain for luminance contrasts for these observers (Fig. 8b, Supplementary Tab. S8). A nested likelihood ratio test, in which the nested model fixed g’across groups, provided no evidence for a difference among the values (*χ*^2^(2) = 2.58, *p* = 0.276) (Supplementary Tab. S16). A test on the individual values did not support the hypothesis that the anomalous values differed from the normal (P vs N: t(429) = 1.40; p = 0.164; D vs N: t(429) = 1.46, p = 0.146) (Supplementary Tab. S11).

**Fig. 7.**
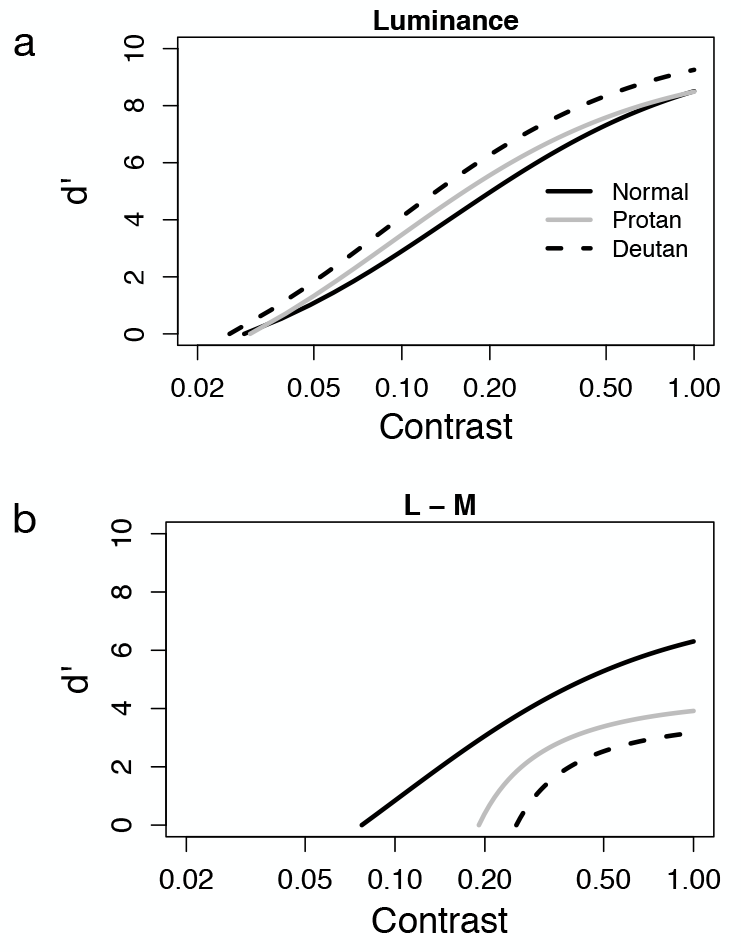
a. Population estimates from nonlinear mixed-effects model for luminance CRDS of Normal (solid black), Protanomalous (solid grey) and Deuteranomalous (dashed black) observers. b. Population estimates from nonlinear mixed-effects model for L-M CRDS in nominal contrast units for the three classes of observers using the same color coding as in a.

Figure 7b shows the population curves for the L-M axis in nominal contrast units. The curves for the anomalous observers asymptote at lower values (Fig. 8a) and rise more steeply (Fig. 8b) than the normal curve (Supplementary Tab. S22). Nested likelihood ratio tests confirmed the differences both for *R_m_* (*χ*^2^(2) = 31.99, p ≪ 0.001) (Supplementary Tab. S27) and for g’ (*χ*^2^(2) = 24.34, p = ≪ 0.001) (Supplementary Tab. S30). The differences in contrast gain along the L-M axis cannot be accounted for by the expression of the contrasts in nominal units as the transformation to cone contrasts only translates the CRDS curves along the log contrast axis without changing their shape.

**Fig. 8.**
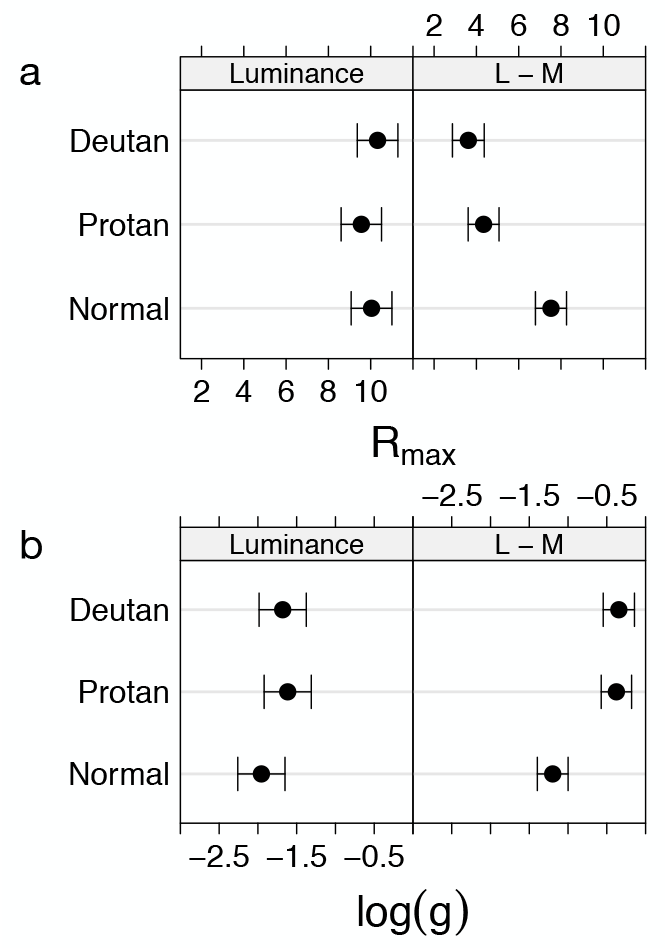
a. Population mean and 95% confidence intervals of the response gain parameter, Rm for normal and anomalous trichromatic observers for CRDSs measured along the luminance and L-M axes. b. Population mean and 95% confidence intervals of the log contrast gain, g’, for normal and anomalous trichromatic observers for CRDSs measured along the luminance and L-M axes.

## 4. Discussion

### 4.1. MLDS

We have demonstrated that MLDS is an effective method for obtaining estimates of the change in appearance of Gabor patterns over a contrast range not accessible with threshold measures of contrast sensitivity. Figure 9 shows a psychometric function for luminance contrast detection (dashed curve) based on the ModelFest data set [38] for estimating threshold of a Gabor stimulus at 1 c/deg using a Weibull function. The solid curve replots the normal luminance MLDS population curve from Fig. 7a. It is notable that there is no stimulus range overlap between the increasing sections of each function. The psychometric function yields information at low contrast values over a four-fold range, whereas the MLDS curve yields information over suprathreshold contrast values spanning a thirty-fold range.

**Fig. 9.**
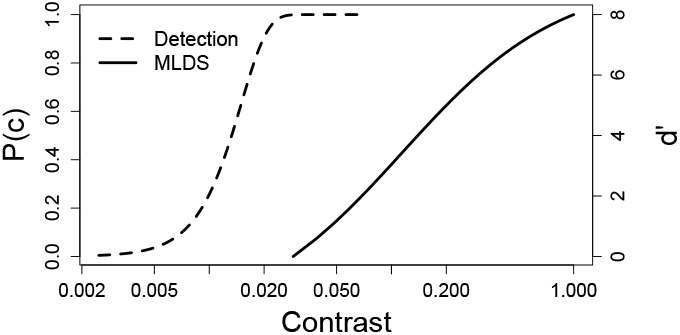
Comparison of contrast, *c*, ranges over which the psychometric function for luminance contrast detection and MLDS scaling operate. The psychometric function (dashed curve and left ordinate values) is a Weibull function, 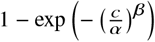, with threshold, *α* based on an estimation of the threshold of a 1 c/deg luminance Gabor function from the ModelFest data set [38] with *β* = 3. The MLDS function (solid curve and right ordinate values) represents the normal population luminance CRDS replotted from Fig. 7a.

The method that we describe is efficient. For 9 contrast levels, as used here, an experienced observer can typically complete the 84-triad session in 2–3 min with data that yield a relatively accurate estimation of the curve. Naive observers require practice sessions to understand the task and stabilize their criteria and generally require longer to complete a session. In both cases, repeated sessions lead to more precise estimates of the curve shape and the fitted parameters.

Contrast response can also be estimated with pedestal experiments. These require estimating a discrimination threshold at each contrast level tested, and the underlying response function is indirectly estimated based on the hypothesis that size of the discrimination threshold is inversely proportional to the underlying response. Such estimates have yielded a slightly more complex functional form to describe the contrast response function in which each term in the numerator and the denominator of the Michaelis-Menten function is raised to a positive exponent [39]. This yields an accelerating response function at low contrasts and a compressive one at high. We found that we could fit our data well assuming that such an exponent is equal to unity. The suprathreshold levels at which MLDS is conducted may not cover the range over which an accelerating nonlinearity is necessary to describe contrast response.

The representation of the MLDS response scale in terms of the signal detection parameter *d′* is based on a parameterization that sets the judgment noise equal to unity on the response scale. How this measure of *d′* relates to that obtained from discrimination experiments is an unsettled question [31, 32] that perhaps can be answered by comparing MLDS response estimates directly with those from discrimination experiments.

Finally, we propose that this approach could be valuable in assessing visual function in pathology of visual pathways. While patients will likely be more challenging to test than observers with normal vision, it would be of interest to identify conditions that differentially influence response and contrast gain and compare the MLDS estimates with those obtained by other non-invasive methods, such as VEPs and functional imagery. In this direction, Crognale et al. [20] reported that VEP amplitudes to luminance contrast saturated near 50% contrast. This value compares favorably with the average MLDS estimate of normal observers along the luminance axis at 50% contrast that is nearly 90% of the predicted response at 100% contrast. Another study used MLDS to scale contrast appearance for square-wave, radial checkerboards and correlated the scales to responses generated using functional cerebral imagery with respect to cortical area V1 and sub-cortical areas LGN and superior colliculus in each of three age groups [40]. As in the current study, these results were well described by Eq. 5. Analyses of the response and contrast gain parameters indicated significant decreases in response gain and increases in contrast gain with age.

### 4.2. Minimum Perceived Contrast, *c*_0_

The stimulus levels used in an MLDS experiment must be above threshold and, in principle, the observer should be capable of sorting them in order [25]. Thus, to obtain measures over the widest contrast range for each observer, we estimated a minimal contrast that could be reliably perceived that we denoted by *c*_0_.

We observed that *c*_0_ did not differ significantly among normal and anomalous observers along the luminance axis. Thus, we do not confirm a recent study reporting that anomalous observers display higher luminance contrast sensitivity than normal observers [41]. This may reflect that *c*_0_, does not, strictly speaking, correspond to a measure of contrast threshold. We did find that anomalous observers, on average, displayed higher luminance contrast gain (Fig. 8a), but the differences observed did not attain statistical significance. We estimate that the number of observers would need to be quadrupled in order to evaluate if the small gain difference observed is reliable.

Expressed in the nominal display contrasts, the *c*_0_ value of anomalous observers along the L-M axis was on average almost three times higher than that of normal observers. This is consistent with the reduced peak-to-trough reduction in chromatic difference signals at the input described in Sect. 1 and in agreement with the findings of Boehm et al. [21] with respect to chromatic discrimination loss in anomalous observers. Nevertheless, when expressed as cone contrasts using average cone fundamentals for normal and anomalous observers, the values converged for all three classes of observers. This supports the hypothesis that normal and anomalous observers require similar response differences to perceive contrast at the physiological level, and is consistent with the analyses of Pokorny and Smith [37] that show that anomalous discrimination can be mapped onto normal discrimination over a reduced stimulus range.

### 4.3. Contrast Appearance along the L-M Axis

Along the L-M axis, anomalous observers showed reduced response gain and increased contrast gain with respect to normal observers (Figs. 7–8). *R_m_* values were, on average, 53% of normal while g values were 229% times the normal value. Recall from Fig. 2 that the effect of reduced spectral separation translates the contrast response curve on a log contrast axis with no change in the steepness of the curves nor in the maximum response. Given the null model based on attenuation of the effective contrast due to reduced spectral separation of the cones, this would require gain compensation in the range of a factor of 6–9.

It is unlikely that the steeper contrast response of anomalous observers along the L-M axis can be explained by luminance artefacts introduced by individual differences from the average luminosity curve. At 1 c/deg, the band-pass luminance contrast sensitivity has diminished to about half of its peak sensitivity [38] while the low-pass L-M chromatic contrast sensitivity remains at its maximal value [42]. This is reflected in the CRDSs when plotted on a cone contrast scale (Figs. 5d–f and Supplementary Figs. 1–3) on which the initial rise of the L-M curves occurs largely below the minimum contrast of the luminance curves.

The steeper rise of *d′* along the contrast axis in Fig. 7b predicts that at contrasts just above threshold, anomalous observers will show enhanced contrast discrimination along the L-M axis. At high contrasts the curves flatten out, indicating that contrast discrimination would become worse, perhaps even showing saturating behavior as in rod discrimination at high luminance levels [43]. These results support MacLeod’s prediction of response in the presence of output noise and Boehm et al.’s conclusions of a post-receptoral gain amplification [21].

Since changes in the terms *α* and *ς* in Eq. 1 do not account for reductions in response gain, some other mechanism must be active. The most parsimonious explanation would require a trade-off in response and contrast gain at the level of the neural mechanism that determines these two parameters. This hypothesis would imply a systematic relation between response and contrast gain above and beyond the effects of the axis tested and the classification of color vision type. Figure 10 shows a scatter plot of the log contrast gain as a function of response gain for all of the observers and conditions tested in this study. The solid gray line corresponds to the best fit linear regression, given by the equation *g′* = 0.36 − 0.21 *R_m_* and indicates a strong linear relation between these two parameters (F(1, 52) = 116.3, p ≪ 0.001; *r*^2^ = 0.69).

**Fig. 10.**
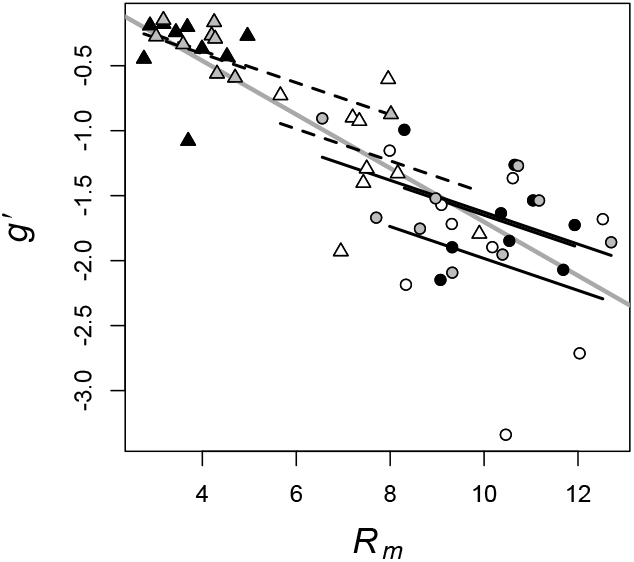
Log contrast gain, *g′*, as a function of response gain, *R_m_*, along luminance (circles) and L-M (triangles) axes, for normal (white), protan (grey) and deutan (black) observers. The grey line is the predicted linear regression for all of the data. The black line segments are the best linear fits to the luminance (solid) and L-M (dashed) values for each of the three types of observers.

It is possible, however, that the relation is carried by the differences between axes and groups and that the linear relation within subcategories is not significant, i.e., demonstrating Simpson’s paradox. To evaluate this possibility, we fit a linear model in which separate regressions were performed within each combination of axis and color vision type. We compared this model with a nested model in which the intercepts varied between groups but the slopes were constrained to be equal. Nested likelihood ratio tests indicated no differences between these models (F(42, 7) = 0.16, p = 0.99). The black solid and dashed line segments in Fig. 10 are the fitted line segments to each combination of axis and color vision type for the constant slope model. The estimated slope from this model is 40% lower than that of the global fit (slope (95% conf. int.): −0.123 (−0.192, −0.054)). Evidence in favor of this model over the global fit (grey line) was obtained from a nested likelihood ratio test (F(49, 3) = 2.86; p = 0.048). Stronger evidence was obtained by performing the same comparisons but introducing an observer specific random intercept in the framework of a linear mixed-effects model (*χ*^2^(3) = 14.05; p = 0.003) [44]. The results support a significant relation between the contrast and response gain independently of the differences due to color axis and color vision type.

### 4.4. Implications for Compensation

Our results suggest a plasticity in visual system organization that optimizes the mapping between stimuli and neural response [23, 24]. Observers with anomalous trichromacy displayed higher contrast gain along a chromatic axis in color space despite a reduced chromatic signal at the input. In the case of these observers, the loss of signal is congenital. There is previous evidence, however, for dynamic changes to the input in normal sensory systems. For example, Kwon et al. [45] exposed normal observers to contrast reduced environments for several hours and observed enhancements in contrast discrimination psychophysically and in neural responses in cortical areas V1 and V2 measured with functional cerebral imagery. Unlike the current results, the model that best described their data required a change in response gain alone and no change in contrast gain.

It has previously been suggested that the loss of chromatic signal in anomalous trichromats due to reduced separation of the cone spectral sensitivities might be ameliorated by the use of selective filters that would act to increase their spectral separation [46, 47]. The MLDS paradigm for measuring contrast appearance would be an ideal paradigm for studying the long-term effects of such aids on contrast perception.

## 5. Conclusion

Contrast response as estimated with a maximum likelihood scaling method is well described by a Michaelis-Menten function. The minimum suprathreshold contrast estimated using the MLDS method was similar for normal and anomalous trichromats along each of the luminance and L-M axes when expressed as cone contrast, suggesting that the neural requirements for detecting contrast are the same for both sets of observers. Anomalous trichromats display a reduced response gain along the L-M axis but do not differ from normal trichromats for luminance contrast. Anomalous trichromats display higher contrast gain than normal observers along the L-M axes. Neither of these differences is predicted by a simple reduction in separation of the cone spectral sensensitivities. It is proposed that the L-M enhancement in contrast gain is due to a post-receptoral gain amplification. A significant linear relation between contrast and response gain across all conditions raises the possibility of a neural trade-off between contrast and response gains.

## Supporting information

Supplemental figures, data tables, and statistical analyses

## Appendix

## Funding

KK was supported by the following grants: LABEX CORTEX (ANR-11-LABX-0042) of Université de Lyon (ANR-11-IDEX-0007) operated by the French National Research Agency (ANR), ANR-15-CE32-0016 CORNET, ANR-17-NEUC-0004, A2P2MC, ANR-17-HBPR-0003, CORTICITY, ANR-19-CE37-0000, DUAL_STREAMS. JSW and BMA were supported by the National Eye Institute (R01 EY 024239).

## Acknowledgments

We thank Susan Garcia for assistance in subject screening and testing. We thank Joshua A. Solomon and Donald I. A. MacLeod for critical comments.

---

**Fig. A1.**
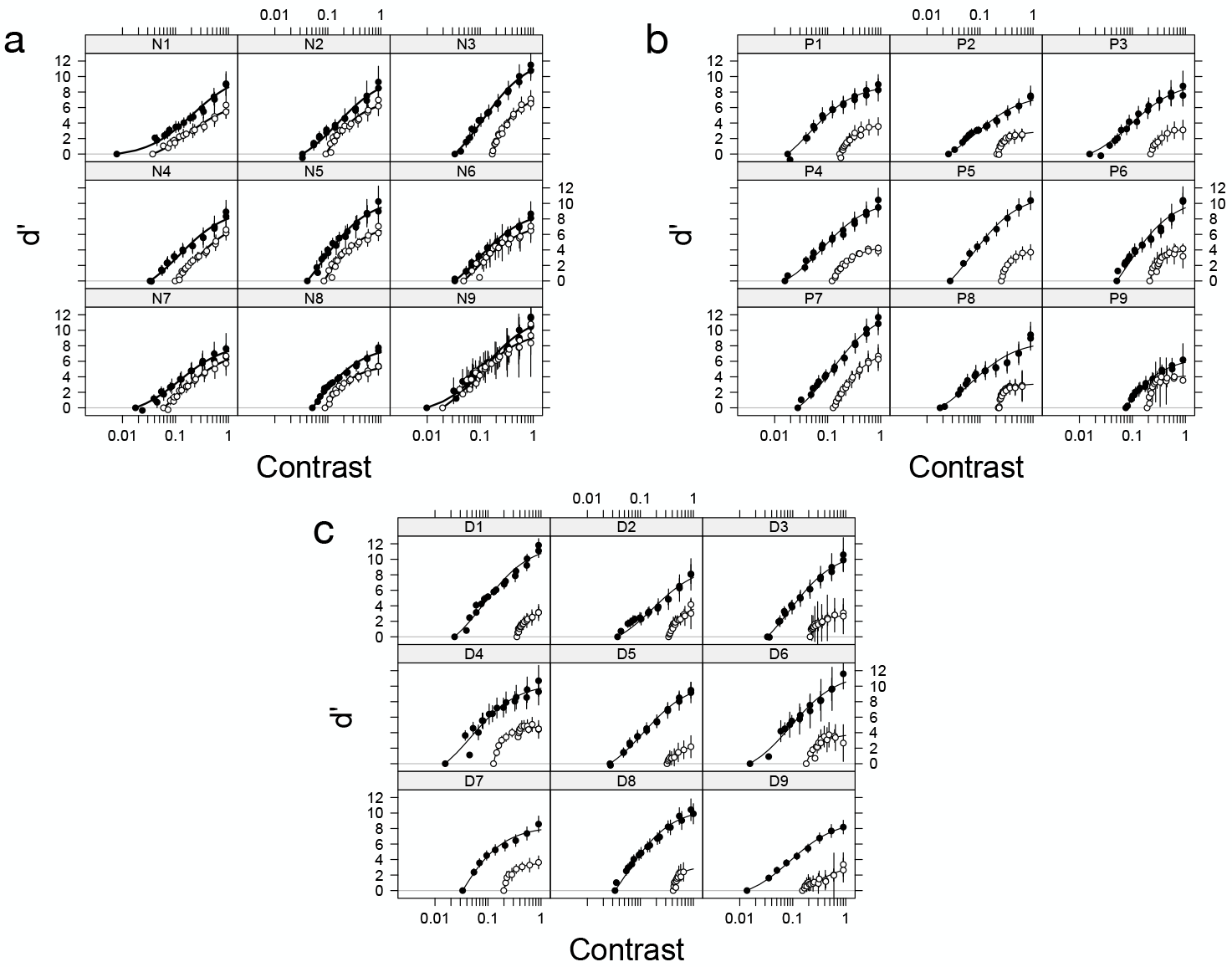
CRDSs parameterized in terms of d’ for all individual observers on a nominal contrast scale, where the maximum contrast value corresponds to the maximum attainable contrast on the display. The black symbols are for measurements along the luminance axis and the white along the L-M axis. The curves are the best-fit Michaelis-Menten functions by a least-squares method. a. Normal, b. Protanomalous, c. Deuteranomalous

## Disclosures

The authors declare no conflicts of interest.

## Notes

https://www.sbri.fr/sites/default/files/supplementary.pdf

