## Supplemental figures, data tables, and statistical analyses for "Suprathreshold contrast response in normal and anomalous trichromats"

In this supplementary section, we present the individual suprathreshold Contrast Response Difference Scales (CRDS) of all observers tested from normal (Fig. S1), protanomalous (Fig. S2) and deuteranomalous (Fig. S3) groups. The data are presented on a scale expressed in terms of cone contrast. Cone contrast, $\mathcal{C}$, for a normal observer was computed from the formula

$$\mathcal{C} = \sqrt{\left(\frac{\Delta L}{L}\right)^2 + \left(\frac{\Delta M}{M}\right)^2 + \left(\frac{\Delta S}{S}\right)^2}, \quad \text{(S1)}$$

where the denominator of each fraction is the cone response to the background and the numerator is the magnitude of the peak deviation of the Gabor stimulus at maximal nominal contrast [1]. Equation S1 was also used for anomalous observers with the anomalous cone responses based on their respective cone spectral sensitivities substituted for the approproriate terms [2]. These values were used to correct the L-M cone contrasts in the figures below, as the luminance contrast is unaffected by the calculation. For stimuli along the L-M axis, the S cone term is constant. Therefore, $\Delta S = 0$ and the S-cones do not contribute to the cone contrast.

In Tabs. S1–S2 , the parameter estimates and associated standard errors of $R_m$, $\varsigma$ and the minimally perceived contrast ($c_0$) for the individually fitted Michaelis-Menten curves in each graph are presented for the luminance and L-M contrast conditions, respectively. In Tabs. S3–S5, the results of the analyses of the $c_0$ values are presented. In Tabs. S6–S30, the results of fitting nonlinear mixed-effect models to the data sets are presented. The fixed-effects and variance components are presented in separate tables. Tables of the results of likelihood ratio tests are presented for pairs of nested models to test for the significance of differences in the parameter estimates.

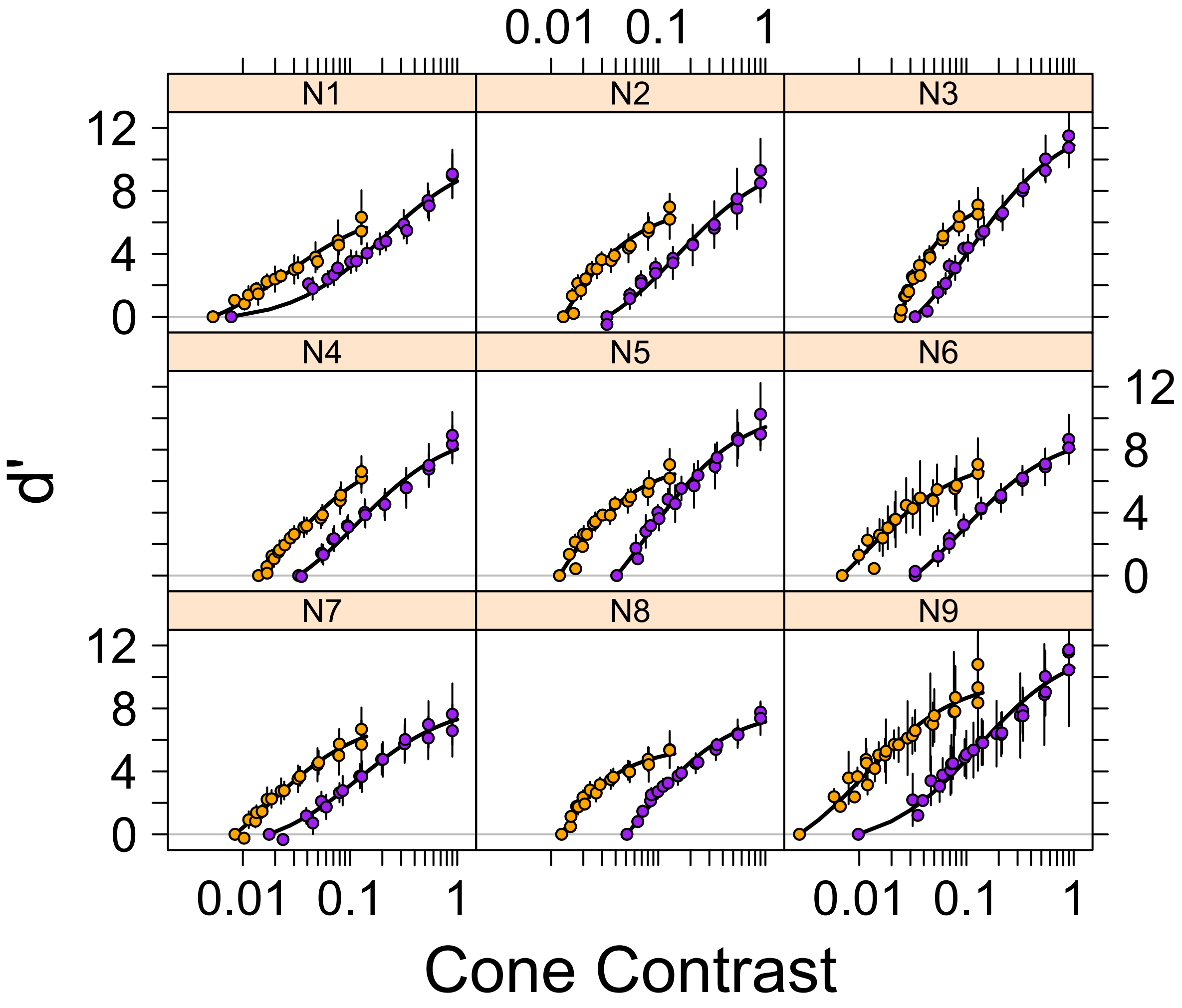


Fig. S1: CRDSs parameterized in terms of $d'$ for individual normal observers on a cone contrast scale. The abscissa shows logarithmic spacing of contrasts to facilitate visualization of the data. The purple symbols are for measurements along the luminance axis and the orange along the L-M axis. The curves are the best-fit Michaelis-Menten functions by a least-squares method.

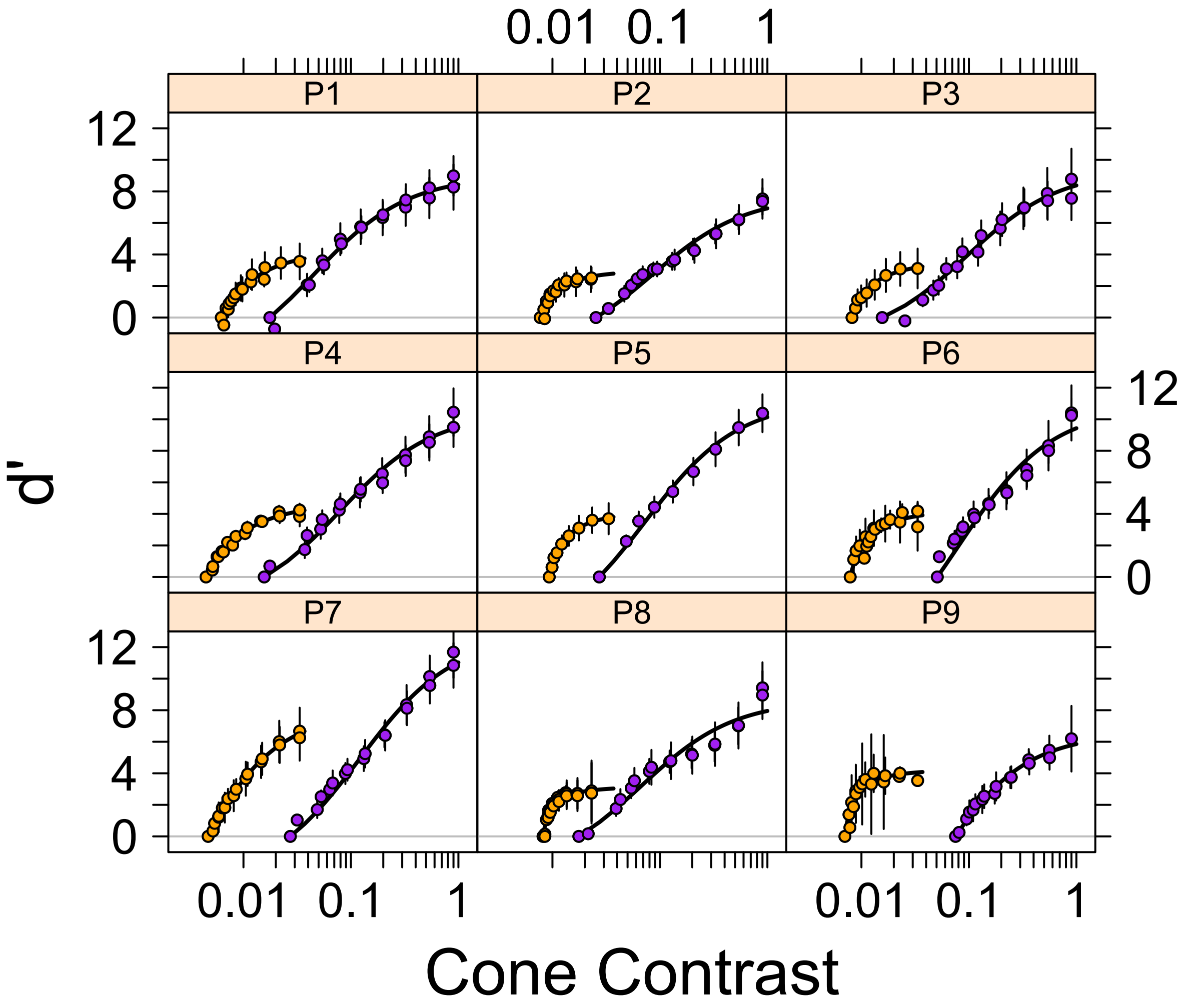


Fig. S2: CRDSs parameterized in terms of $d'$ for individual protanomalous observers on a cone contrast scale. The abscissa shows logarithmic spacing of contrasts to facilitate visualization of the data. The purple symbols are for measurements along the luminance axis and the orange along the L-M axis. The curves are the best-fit Michaelis-Menten functions by a least-squares method.

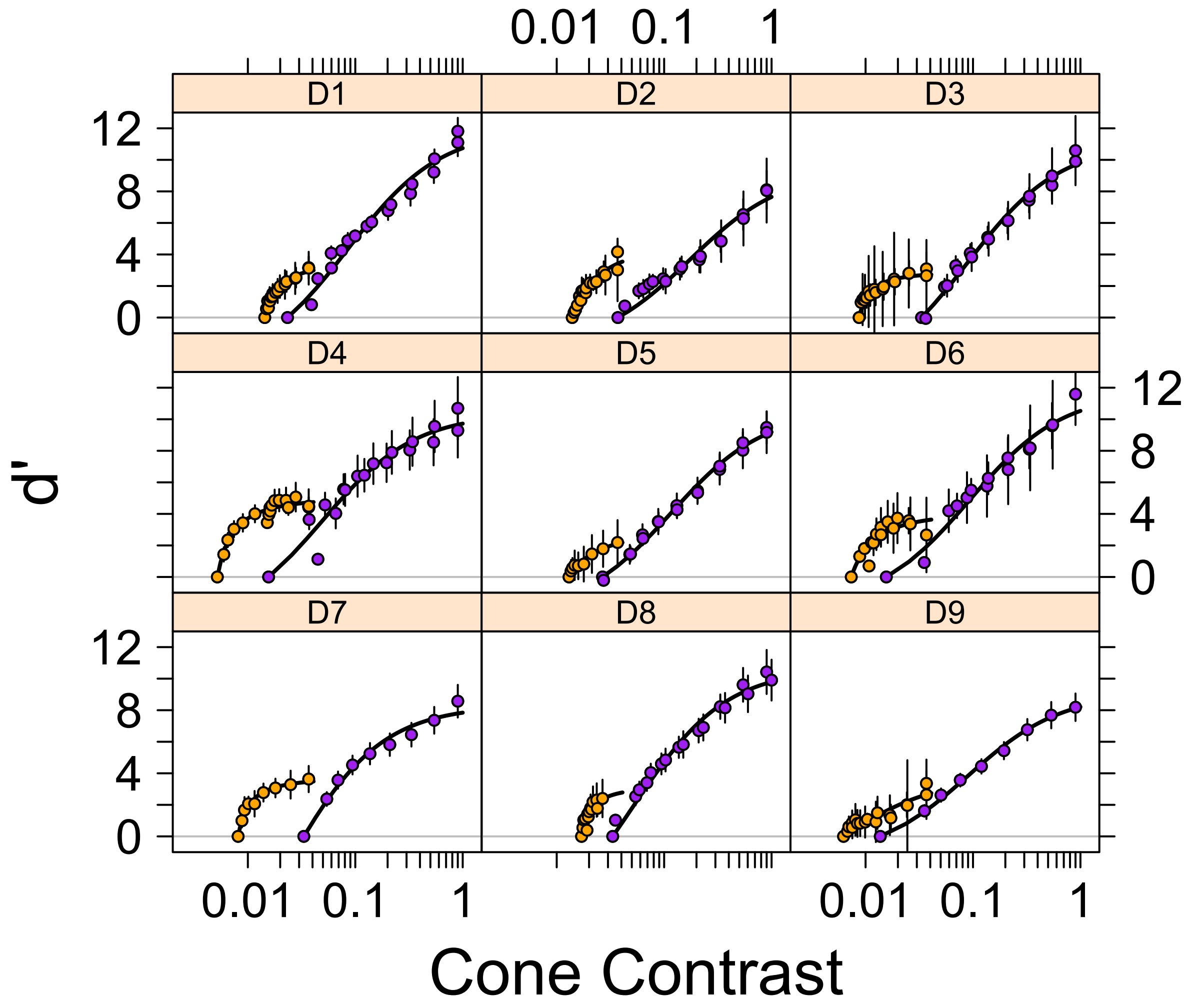


Fig. S3: CRDSs parameterized in terms of $d'$ for individual deuteranomalous observers on a cone contrast scale. The abscissa shows logarithmic spacing of contrasts to facilitate visualization of the data. The purple symbols are for measurements along the luminance axis and the orange along the L-M axis. The curves are the best-fit Michaelis-Menten functions by a least-squares method.

#### Parameter estimates for Michaelis-Menten fits to luminance CRDSs by individual

| ID | Type | $R_m$ | $SE_{R_m}$ | $\varsigma$ | $SE_\varsigma$ | $c_0$ | Gain |
|---|---|---|---|---|---|---|---|
| N1 | Normal | 10.462 | 0.525 | 0.212 | 0.027 | 0.008 | 0.035 |
| N2 | Normal | 10.176 | 0.532 | 0.188 | 0.026 | 0.033 | 0.150 |
| N3 | Normal | 12.528 | 0.388 | 0.145 | 0.013 | 0.033 | 0.186 |
| N4 | Normal | 9.312 | 0.464 | 0.152 | 0.021 | 0.033 | 0.179 |
| N5 | Normal | 10.611 | 0.378 | 0.120 | 0.013 | 0.041 | 0.255 |
| N6 | Normal | 9.091 | 0.264 | 0.127 | 0.011 | 0.033 | 0.208 |
| N7 | Normal | 8.335 | 0.329 | 0.139 | 0.016 | 0.018 | 0.112 |
| N8 | Normal | 7.989 | 0.271 | 0.111 | 0.011 | 0.051 | 0.316 |
| N9 | Normal | 12.034 | 0.454 | 0.138 | 0.015 | 0.010 | 0.066 |
| D1 | Deutan | 11.929 | 0.491 | 0.108 | 0.014 | 0.023 | 0.178 |
| D2 | Deutan | 9.319 | 0.678 | 0.210 | 0.039 | 0.037 | 0.150 |
| D3 | Deutan | 11.043 | 0.355 | 0.121 | 0.012 | 0.033 | 0.215 |
| D4 | Deutan | 10.360 | 0.469 | 0.064 | 0.010 | 0.016 | 0.195 |
| D5 | Deutan | 10.538 | 0.290 | 0.143 | 0.011 | 0.027 | 0.158 |
| D6 | Deutan | 11.686 | 0.573 | 0.108 | 0.016 | 0.016 | 0.126 |
| D7 | Deutan | 8.301 | 0.385 | 0.056 | 0.010 | 0.033 | 0.370 |
| D8 | Deutan | 10.649 | 0.300 | 0.084 | 0.008 | 0.033 | 0.283 |
| D9 | Deutan | 9.071 | 0.210 | 0.104 | 0.008 | 0.014 | 0.117 |
| P1 | Protan | 8.968 | 0.244 | 0.063 | 0.006 | 0.018 | 0.219 |
| P2 | Protan | 7.703 | 0.320 | 0.109 | 0.014 | 0.025 | 0.188 |
| P3 | Protan | 9.323 | 0.350 | 0.111 | 0.013 | 0.016 | 0.123 |
| P4 | Protan | 10.393 | 0.351 | 0.094 | 0.010 | 0.016 | 0.142 |
| P5 | Protan | 11.171 | 0.424 | 0.100 | 0.012 | 0.027 | 0.215 |
| P6 | Protan | 10.723 | 0.660 | 0.130 | 0.023 | 0.051 | 0.281 |
| P7 | Protan | 12.706 | 0.445 | 0.148 | 0.015 | 0.027 | 0.156 |
| P8 | Protan | 8.634 | 0.501 | 0.084 | 0.016 | 0.018 | 0.173 |
| P9 | Protan | 6.554 | 0.226 | 0.111 | 0.011 | 0.075 | 0.404 |

Table S1: Michaelis-Menten parameter estimates, $R_m$ and $\varsigma$, for individual observers for luminance contrast. Standard errors (SE) were obtained from the variance-covariance matrix at the maximum likelihood parameter estimates. $c_0$ is the minimum perceived contrast estimate. Gain is defined as $c_0/(c_0 + \varsigma)$.

### Parameter estimates for Michaelis-Menten fits to L–M CRDSs by individual

| ID | Type | $R_m$ | $SE_{R_m}$ | $\varsigma$ | $SE_\varsigma$ | $c_0$ | Gain |
|---|---|---|---|---|---|---|---|
| N1 | Normal | 6.949 | 0.416 | 0.218 | 0.032 | 0.037 | 0.145 |
| N2 | Normal | 7.201 | 0.401 | 0.133 | 0.021 | 0.092 | 0.407 |
| N3 | Normal | 7.958 | 0.309 | 0.140 | 0.015 | 0.169 | 0.547 |
| N4 | Normal | 8.161 | 0.452 | 0.274 | 0.034 | 0.099 | 0.265 |
| N5 | Normal | 7.344 | 0.400 | 0.129 | 0.020 | 0.084 | 0.396 |
| N6 | Normal | 7.500 | 0.454 | 0.129 | 0.022 | 0.049 | 0.275 |
| N7 | Normal | 7.429 | 0.359 | 0.182 | 0.023 | 0.059 | 0.246 |
| N8 | Normal | 5.662 | 0.196 | 0.095 | 0.010 | 0.088 | 0.483 |
| N9 | Normal | 9.899 | 0.371 | 0.098 | 0.011 | 0.019 | 0.166 |
| D1 | Deutan | 3.437 | 0.163 | 0.096 | 0.012 | 0.351 | 0.785 |
| D2 | Deutan | 4.524 | 0.406 | 0.182 | 0.036 | 0.341 | 0.652 |
| D3 | Deutan | 2.885 | 0.151 | 0.045 | 0.009 | 0.213 | 0.826 |
| D4 | Deutan | 4.961 | 0.161 | 0.039 | 0.007 | 0.127 | 0.763 |
| D5 | Deutan | 2.755 | 0.390 | 0.179 | 0.060 | 0.318 | 0.640 |
| D6 | Deutan | 3.992 | 0.486 | 0.080 | 0.032 | 0.180 | 0.692 |
| D7 | Deutan | 3.680 | 0.153 | 0.045 | 0.007 | 0.198 | 0.816 |
| D8 | Deutan | 3.172 | 0.636 | 0.081 | 0.037 | 0.416 | 0.836 |
| D9 | Deutan | 3.698 | 0.665 | 0.296 | 0.115 | 0.152 | 0.340 |
| P1 | Protan | 4.312 | 0.346 | 0.127 | 0.025 | 0.168 | 0.570 |
| P2 | Protan | 3.016 | 0.235 | 0.066 | 0.015 | 0.208 | 0.760 |
| P3 | Protan | 3.590 | 0.167 | 0.088 | 0.013 | 0.221 | 0.715 |
| P4 | Protan | 4.697 | 0.136 | 0.098 | 0.009 | 0.121 | 0.553 |
| P5 | Protan | 4.197 | 0.093 | 0.077 | 0.005 | 0.252 | 0.765 |
| P6 | Protan | 4.272 | 0.301 | 0.072 | 0.017 | 0.212 | 0.747 |
| P7 | Protan | 8.016 | 0.168 | 0.177 | 0.010 | 0.127 | 0.417 |
| P8 | Protan | 3.170 | 0.124 | 0.035 | 0.005 | 0.221 | 0.862 |
| P9 | Protan | 4.252 | 0.238 | 0.034 | 0.008 | 0.190 | 0.848 |

Table S2: Michaelis-Menten parameter estimates, $R_m$ and $\varsigma$, for individual observers for L-M contrast. Standard errors (SE) were obtained from the variance-covariance matrix at the maximum likelihood parameter estimates. $c_0$ is the minimum perceived contrast estimate. Gain is defined as $c_0/(c_0 + \varsigma)$.

#### Analysis of $c_0$ values

| | Df | Sum Sq | Mean Sq | F value | Pr(>F) |
|---|---|---|---|---|---|
| Axis | 2 | 470.195 | 235.097 | 927.073 | $p \ll 0.001$ |
| Axis:Type | 4 | 8.292 | 2.073 | 8.175 | $p \ll 0.001$ |
| Residuals | 48 | 12.172 | 0.254 | | |

Table S3: Analysis of variance table for modeling the minimum perceived contrast, $\log_e(c_0)$, by factors Axis with levels luminance and L-M and Type with levels normal, protanomalous and deuteranomalous. In the model, Type was nested within the factor Axis so that the interaction term above and the coefficients in Tab. S4 indicate the difference between groups along a given axis.

| | Estimate | Std. Error | t value | Pr(>\|t\|) |
|---|---|---|---|---|
| Lum | -3.701 | 0.168 | -22.049 | $p \ll 0.001$ |
| L-M | -2.717 | 0.168 | -16.189 | $p \ll 0.001$ |
| Lum(D) - Lum(N) | -0.021 | 0.237 | -0.089 | 0.930 |
| L-M(D) - L-M(N) | 1.276 | 0.237 | 5.374 | $p \ll 0.001$ |
| Lum(P) ) - Lum(N) | 0.052 | 0.237 | 0.218 | 0.828 |
| L-M(P) - L-M(N) | 1.035 | 0.237 | 4.358 | $p \ll 0.001$ |

Table S4: Summary results of the coefficients for the terms in the two-way ANOVA fit to the $\log_e(c_0)$ values. The Lum and L-M terms correspond to the mean estimates for the normal group along the luminance and L-M axes, respectively. The subsequent terms indicate differences of each anomalous group from the normal estimates, treated as controls, so that the p-values indicate the probability of the difference observed under the null hypothesis of no difference from the normal group.

| | Df | Sum Sq | Mean Sq | F value | Pr(>F) |
|---|---|---|---|---|---|
| Type | 2 | 0.640 | 0.320 | 1.497 | 0.244 |
| Residuals | 24 | 5.134 | 0.214 | | |

Table S5: One-way analysis of variance comparing the means across the three groups along the L-M axis with the nominal contrasts expressed as cone contrasts.

### Luminance Contrast Analyses

| | Estimate | Std.Error | DF | t-value | p-value |
|---|---|---|---|---|---|
| $R_m$(N) | 10.051 | 0.496 | 397.000 | 20.283 | p $\ll$ 0.001 |
| $R_m$(P) - $R_m$(N) | -0.483 | 0.763 | 397.000 | -0.633 | 0.527 |
| $R_m$(D) - $R_m$(N) | 0.279 | 0.624 | 397.000 | 0.447 | 0.655 |
| $g'$(N) | -1.958 | 0.218 | 397.000 | -8.993 | p $\ll$ 0.001 |
| $g'$(P) - $g'$(N) | 0.343 | 0.244 | 397.000 | 1.403 | 0.161 |
| $g'$(D) - $g$('N) | 0.276 | 0.246 | 397.000 | 1.122 | 0.263 |

Table S6: Fixed-effects results for testing luminance CRDS from a nonlinear mixed-effects model fit to the data in which the two parameters, $R_m$ and $g' = \log(g)$, and their variances were free to vary for each group. The Estimate is the least-squares population estimate. The standard errors are obtained from the variance-covariance matrix at the maximum likelihood solution. The succeeding columns show the calculation and p-values for the Wald statistics, i.e., the ratio of the Estimate and its standard error, treated as a t-distributed statistic because the variance is estimated from the data independently of the mean. The model is fit with treatement contrasts so that the base level is the estimate for the normal (N) group and the succeeding levels indicate contrasts with this level for the protanomalous (P) and deuteranomlous (D) groups, respectively.

| | Variance | StdDev | Corr | | | | |
|---|---|---|---|---|---|---|---|
| $R_m$(N) | 1.967 | 1.402 | $R_m$(N) | $R_m$(P) | $R_m$(D) | $g'$(N) | $g'$(P) |
| $R_m$(P) | 0.927 | 0.963 | -0.030 | | | | |
| $R_m$(D) | 0.903 | 0.950 | -0.672 | 0.027 | | | |
| $g'$(N) | 0.408 | 0.639 | -0.308 | 0.011 | 0.289 | | |
| $g'$(P) | 0.176 | 0.419 | 0.203 | -0.109 | -0.224 | -0.909 | |
| $g'$(D) | 0.165 | 0.406 | 0.266 | -0.023 | -0.289 | -0.900 | 0.812 |
| Residual | 0.259 | 0.509 | | | | | |

Table S7: Variance components of the observer dependent random effects for $R_m$ and $g'$ for the luminance CRDS from a nonlinear mixed-effects model fit to the data in which the parameters for each group and their variances were free to vary. The StdDev column is the square root of the Variance column and provides a value more easily interpretable on the scale of the data. The Corr columns show the correlations between the variance estimates of the two parameters. The Residual row indicates the variance unaccounted for by the variances associated with the two random effects.

| | Estimate | Std.Error | DF | t-value | p-value |
|---|---|---|---|---|---|
| $R_m$(N) | 10.041 | 0.494 | 397.000 | 20.337 | p $\ll$ 0.001 |
| $R_m$(P) - $R_m$(N) | -0.483 | 0.695 | 397.000 | -0.695 | 0.487 |
| $R_m$(D) - $R_m$(N) | 0.284 | 0.697 | 397.000 | 0.408 | 0.684 |
| $g'$(N) | -1.955 | 0.156 | 397.000 | -12.543 | p $\ll$ 0.001 |
| $g'$(P) - $g'$(N) | 0.339 | 0.220 | 397.000 | 1.539 | 0.125 |
| $g'$(D) - $g$('N) | 0.275 | 0.220 | 397.000 | 1.246 | 0.214 |

Table S8: Fixed-effects results for testing luminance CRDS from a nonlinear mixed-effects model fit to the data in which the two parameters, $R_m$ and $g' = \log(g)$, were free to vary for each group. The Estimate is the least-squares population estimate. The standard errors are obtained from the variance-covariance matrix at the maximum likelihood solution. The succeeding columns show the calculation and p-values for the Wald statistics, i.e., the ratio of the Estimate and its standard error, treated as a t-distributed statistic because the variance is estimated from the data independently of the mean. The model is fit with treatement contrasts so that the base level is the estimate for the normal (N) group and the succeeding levels indicate contrasts with this level for the protanomalous (P) and deuteranomlous (D) groups, respectively.

| | Variance | StdDev | Corr |
|---|---|---|---|
| $R_m$ | 1.953 | 1.397 | |
| $g'$ | 0.203 | 0.451 | -0.276 |
| Residual | 0.259 | 0.509 | |

Table S9: Variance components of the observer dependent random effects for $R_m$ and $g'$ for the luminance CRDS from a nonlinear mixed-effects model fit to the data in which the parameters for each group were free to vary. The StdDev column is the square root of the Variance column and provides a value more easily interpretable on the scale of the data. The Corr column shows the correlation between the variance estimates of the two parameters. The Residual row indicates the variance unaccounted for by the variances associated with the two random effects.

| | Model | df | logLik | Test | $\chi^2$ | p-value |
|---|---|---|---|---|---|---|
| Variances fixed in-group | 1 | 10 | -408.045 | | | |
| Variances not-fixed in-group | 2 | 28 | -404.277 | 1 vs 2 | 7.537 | 0.985 |

Table S10: Nested likelihood ratio test for comparison between models in which the variances of the parameters were free to vary between groups and in which they were fixed for the three groups.

| | Estimate | Std.Error | DF | t-value | p-value |
|---|---|---|---|---|---|
| $R_m$ | 9.973 | 0.290 | 399.000 | 34.360 | p $\ll$ 0.001 |
| $g'$(N) | -1.948 | 0.150 | 399.000 | -12.984 | p $\ll$ 0.001 |
| $g'$(P) - $g'$(N) | 0.291 | 0.208 | 399.000 | 1.395 | 0.164 |
| $g'$(D) - $g'$(N) | 0.303 | 0.208 | 399.000 | 1.455 | 0.146 |

Table S11: Fixed-effects results for testing luminance CRDSs from a nonlinear mixed-effects model in which $R_m$ was constrained to be the same for the three groups of observers.

| | Variance | StdDev | Corr |
|---|---|---|---|
| $R_m$ | 2.060 | 1.435 | |
| $g'$ | 0.204 | 0.451 | -0.282 |
| Residual | 0.259 | 0.509 | |

Table S12: Variance components of the observer dependent random effects for $R_m$ and $\varsigma$ for the luminance CRDSs from a nonlinear mixed-effects model in which the parameter $R_m$ was constrained to be the same for the three groups of observers.

| | Model | df | logLik | Test | $\chi^2$ | p-value |
|---|---|---|---|---|---|---|
| 1. $R_m$ fixed | 1 | 8 | -408.659 | | | |
| 2. $R_m$ variable | 2 | 10 | -408.045 | 1 vs 2 | 1.228 | 0.541 |

Table S13: Nested likelihood ratio test for comparison of model in which $R_m$ varies across groups with one in which $R_m$ was constrained to be identical across groups.

| | Estimate | Std.Error | DF | t-value | p-value |
|---|---|---|---|---|---|
| $R_m$(N) | 9.833 | 0.476 | 399.000 | 20.654 | p $\ll$ 0.001 |
| $R_m$(P) - $R_m$(N) | -0.141 | 0.658 | 399.000 | -0.214 | 0.831 |
| $R_m$(D) - $R_m$(N) | 0.565 | 0.659 | 399.000 | 0.857 | 0.392 |
| $g'$ | -1.751 | 0.094 | 399.000 | -18.603 | p $\ll$ 0.001 |

Table S14: Fixed-effects results for testing luminance CRDSs from a nonlinear mixed-effects model in which $g'$ was constrained to be the same for the three groups of observers.

| | Variance | StdDev | Corr |
|---|---|---|---|
| $R_m$ | 1.974 | 1.405 | |
| $g'$ | 0.225 | 0.474 | -0.293 |
| Residual | 0.259 | 0.509 | |

Table S15: Variance components of the observer dependent random effects for $R_m$ and $g'$ for the luminance CRDSs from a nonlinear mixed-effects model in which the parameter $g'$ was constrained to be the same for the three groups of observers.

| | Model | df | logLik | Test | $\chi^2$ | p-value |
|---|---|---|---|---|---|---|
| 1. $g'$ fixed | 1 | 8 | -409.334 | | | |
| 2. $g'$ variable | 2 | 10 | -408.045 | 1 vs 2 | 2.578 | 0.276 |

Table S16: Nested likelihood ratio test for comparison of model in which $g'$ varies across groups with one in which $g'$ was constrained to be identical across groups.

| | Estimate | Std.Error | DF | t-value | p-value |
|---|---|---|---|---|---|
| $R_m$ | 9.974 | 0.290 | 401.000 | 34.448 | p $\ll$ 0.001 |
| $g'$ | -1.750 | 0.094 | 401.000 | -18.667 | p $\ll$ 0.001 |

Table S17: Fixed-effect estimates from a model fit to luminance CRDSs in which both $R_m$ and $g'$ were constrained to be equal across groups.

| | Variance | StdDev | Corr |
|---|---|---|---|
| $R_m$ | 2.060 | 1.435 | |
| $g'$ | 0.224 | 0.474 | -0.275 |
| Residual | 0.259 | 0.509 | |

Table S18: Variance components of the observer dependent random effects for $R_m$ and $g'$ for the luminance CRDSs for which the parameters $R_m$ and $g'$ were constrained to be the same for the three groups of observers

| | Model | df | logLik | Test | $\chi^2$ | p-value |
|---|---|---|---|---|---|---|
| 1. $R_m$ and $g'$ fixed | 1 | 6 | -409.959 | | | |
| 2. $R_m$ and $g'$ variable | 2 | 10 | -408.045 | 1 vs 2 | 3.828 | 0.430 |

Table S19: Nested likelihood ratio test for comparison of a model in which both $R_m$ and $g'$ varied across groups with one in which both were constrained to be identical across groups.

### L–M Contrast Analyses

| | Estimate | Std.Error | DF | t-value | p-value |
|---|---|---|---|---|---|
| $R_m$(N) | 7.547 | 0.374 | 389.000 | 20.181 | p $\ll$ 0.001 |
| $R_m$(P) - $R_m$(N) | -3.227 | 0.598 | 389.000 | -5.392 | p $\ll$ 0.001 |
| $R_m$(D) - $R_m$(N) | -3.953 | 0.463 | 389.000 | -8.533 | p $\ll$ 0.001 |
| $g'$(N) | -1.206 | 0.147 | 389.000 | -8.215 | p $\ll$ 0.001 |
| $g'$(P) - $g'$(N) | 0.838 | 0.165 | 389.000 | 5.068 | p $\ll$ 0.001 |
| $g'$(D) - $g'$(N) | 0.889 | 0.153 | 389.000 | 5.813 | p $\ll$ 0.001 |

Table S20: Fixed-effects results for testing L-M CRDSs in a nonlinear mixed-effects model in which the parameters for each group and their variances were free to vary. All other details as in Tab. S6

| | Variance | StdDev | Corr | | | | |
|---|---|---|---|---|---|---|---|
| $R_m$(N) | 1.119 | 1.058 | $R_m$(N) | $R_m$(P) | $R_m$(D) | $g'$(N) | $g'$(P) |
| $R_m$(P) - $R_m$(N) | 0.665 | 0.815 | 0.032 | | | | |
| $R_m$(D) - $R_m$(N) | 0.298 | 0.546 | -0.765 | -0.011 | | | |
| $g'$(N) | 0.184 | 0.429 | -0.439 | -0.299 | 0.453 | | |
| $g'$(P) - $g'$(N) | 0.087 | 0.295 | 0.129 | 0.046 | -0.270 | -0.885 | |
| $g'$(D) - $g'$(N) | 0.261 | 0.511 | 0.458 | 0.312 | -0.459 | -0.992 | 0.860 |
| Residual | 0.127 | 0.357 | | | | | |

Table S21: Variance components of the observer dependent random effects for $R_m$ and $g'$ for the L-M CRDSs in a nonlinear mixed-effects model in which the parameters for each group and their variances were free to vary. All other details as in Tab. S7

| | Estimate | Std.Error | DF | t-value | p-value |
|---|---|---|---|---|---|
| $R_m$(N) | 7.527 | 0.377 | 389.000 | 19.975 | p $\ll$ 0.001 |
| $R_m$(P) - $R_m$(N) | -3.187 | 0.531 | 389.000 | -6.002 | p $\ll$ 0.001 |
| $R_m$(D) - $R_m$(N) | -3.909 | 0.539 | 389.000 | -7.258 | p $\ll$ 0.001 |
| $g'$(N) | -1.198 | 0.102 | 389.000 | -11.780 | p $\ll$ 0.001 |
| $g'$(P) - $g'$(N) | 0.821 | 0.143 | 389.000 | 5.750 | p $\ll$ 0.001 |
| $g'$(D) - $g'$(N) | 0.853 | 0.145 | 389.000 | 5.886 | p $\ll$ 0.001 |

Table S22: Fixed-effects results for testing L-M CRDSs in a nonlinear mixed-effects model in which the parameters for each group were free to vary. All other details as in Tab. S8

| | Variance | StdDev | Corr |
|---|---|---|---|
| $R_m$ | 1.141 | 1.068 | |
| $g'$ | 0.084 | 0.291 | -0.448 |
| Residual | 0.126 | 0.355 | |

Table S23: Variance components of the observer dependent random effects for $R_m$ and $g'$ for the L-M CRDSs in a nonlinear mixed-effects model in which the parameters for each group were free to vary. All other details as in Tab. S9

| | Model | df | logLik | Test | $\chi^2$ | p-value |
|---|---|---|---|---|---|---|
| Variances fixed in-group | 1 | 10 | -249.321 | | | |
| Variances not-fixed in-group | 2 | 28 | -239.930 | 1 vs 2 | 18.782 | 0.405 |

Table S24: Nested likelihood ratio test for comparison between models in which the variances of the parameters were free to vary between groups and in which they were fixed for the three groups.

| | Estimate | Std.Error | DF | t-value | p-value |
|---|---|---|---|---|---|
| $R_m$ | 5.162 | 0.393 | 391.000 | 13.134 | $p \ll 0.001$ |
| $g'$(N) | -0.896 | 0.101 | 391.000 | -8.863 | $p \ll 0.001$ |
| $g'$(P) - $g'$(N) | 0.416 | 0.124 | 391.000 | 3.358 | 0.001 |
| $g'$(D) - $g'$(N) | 0.358 | 0.125 | 391.000 | 2.865 | 0.004 |

Table S25: Fixed-effects results for testing L-M CRDSs from a nonlinear mixed-effects model in which the estimate of $R_m$ was constrained to be equal across groups.

| | Variance | StdDev | Corr |
|---|---|---|---|
| $R_m$ | 3.992 | 1.998 | |
| $g'$ | 0.128 | 0.357 | -0.694 |
| Residual | 0.127 | 0.356 | |

Table S26: Variance components of the observer dependent random effects for $R_m$ and $g'$ for the L-M CRDSs from a nonlinear mixed-effects model in which the estimate of $R_m$ was constrained to be equal across groups. All other details as in Tab. S12

| | Model | df | logLik | Test | $\chi^2$ | p-value |
|---|---|---|---|---|---|---|
| 1. $R_m$ fixed | 1 | 8 | -265.317 | | | |
| 2. $R_m$ variable | 2 | 10 | -249.321 | 1 vs 2 | 31.992 | $p \ll 0.001$ |

Table S27: Likelihood ratio test results comparing models in which $R_m$ was constrained to be fixed or was free to vary across groups, for stimuli varying in contrast along the L-M axis.

| | Estimate | Std.Error | DF | t-value | p-value |
|---|---|---|---|---|---|
| $R_m$(N) | 6.635 | 0.363 | 391.000 | 18.301 | $p \ll 0.001$ |
| $R_m$(P) - $R_m$(N) | 1.833 | 0.459 | 391.000 | -3.993 | $p \ll 0.001$ |
| $R_m$(D) - $R_m$(N) | -2.467 | 0.465 | 391.000 | -5.302 | $p \ll 0.001$ |
| $g'$ | -0.656 | 0.098 | 391.000 | -6.698 | $p \ll 0.001$ |

Table S28: Fixed-effects results for testing L-M CRDSs from a nonlinear mixed-effects model in which the estimate of $g'$ was constrained to be equal across groups, but not $R_m$.

| | Variance | StdDev | Corr |
|---|---|---|---|
| $R_m$ | 1.514 | 1.231 | |
| $g'$ | 0.247 | 0.497 | -0.644 |
| Residual | 126 | 0.355 | |

Table S29: Variance components of the observer dependent random effects for $R_m$ and $g'$ for the L-M CRDSs in a nonlinear mixed-effects model in which the estimate of $g'$ was constrained to be equal across groups, but not $R_m$.

| | Model | df | logLik | Test | $\chi^2$ | p-value |
|---|---|---|---|---|---|---|
| 1. $g'$ fixed | 1 | 8 | -261.489 | | | |
| 2. $g'$ variable | 2 | 10 | -249.321 | 1 vs 2 | 24.337 | $p \ll 0.001$ |

Table S30: Likelihood ratio test results comparing models in which $g'$ was constrained to be fixed or was free to vary across groups, for stimuli varying in contrast along the L-M axis.
